# CD74^+^ fibroblasts proliferate upon mechanical stretching to promote angiogenesis in keloid

**DOI:** 10.1101/2024.05.05.592627

**Authors:** Jingheng Zhang, Shuyao Li, Chunmei Kuang, Yunfan Shen, Haibin Yu, Fang Chen, Ruijun Tang, Song Mao, Lu Lv, Min Qi, Jianglin Zhang, Kai Yuan

**Author notes:** Correspondence: Kai Yuan,; Jianglin Zhang,; Min Qi. **Author Contributions:** J.H.Z, K.Y, J.L.Z, and M.Q. designed research; J.H.Z and S.Y.L performed the experiment; F.C., R.J.T., S.M. and L.L. analyzed data; J.H.Z, S.Y.L., C.M.K., Y.F.S., H.B.Y, K.Y, J.L.Z, and M.Q wrote the manuscript. **Competing Interests:** The authors declare no competing interest.

## Abstract

The healing of human skin wounds is susceptible to perturbation caused by excessive mechanical stretching, resulting in enlarged scars, hypertrophic scars, or even keloids in predisposed individuals. Keloids are fibro-proliferative scar tissues that extend beyond the initial wound boundary, consisting of the actively progressing leading edge and the quiescent center. The stretch-associated outgrowth and enhanced angiogenesis are two features of the leading edge of keloids. However, which cell population is responsible for transducing the mechanical stimulation to the pathological alterations of keloid tissues remains unclear. Herein, through joint analysis of single-cell RNA sequencing of keloid specimens and RNA sequencing of stretched keloid fibroblasts, we identified CD74^+^ fibroblasts, a previously unappreciated subset of fibroblasts, as a key player in stretch-induced keloid progression. Examination of macrophage markers suggested a possible myeloid origin of the CD74^+^ fibroblasts. Immunostaining of keloid cryosections depicted a predominant distribution of CD74^+^ fibroblasts in the leading edge, interacting with vasculature. CD74^+^ fibroblasts possessed pro-angiogenic and migratory capacities, as revealed by *in vitro* transwell and tube formation assays on purified CD74^+^ fibroblasts. Additionally, these cells underwent proliferation upon stretching, through PIEZO1-mediated calcium influx and the downstream ERK and AKT signaling. Collectively, our findings propose a model wherein CD74^+^ fibroblasts serve as pivotal drivers of stretch-induced keloid progression, fueled by their proliferative, pro-angiogenic, and migratory capacities. Targeting the attributes of CD74^+^ fibroblasts hold promise as a therapeutic strategy for keloid management.

**Significance statement:** Keloids are fibro-proliferative scars resulting from aberrant skin wound healing processes, consisting of the actively progressing leading edge and the quiescent center. Mechanical stretching and neo-vascularization have both been implicated in keloid progression, yet little is known about whether they are interconnected. Herein, we demonstrated that CD74^+^ fibroblasts, a previously undiscovered fibroblast subset, possessed heightened pro-angiogenic and migratory capacities, and underwent proliferation upon mechanical stretching, thereby facilitating the progression of the leading edge of keloids. Examination of macrophage markers suggested a possible myeloid origin of CD74^+^ fibroblasts. Our findings uncover the connection between stretch-induced keloid progression and neo-vascularization through CD74^+^ fibroblasts and provide valuable insights into potential therapeutic interventions.

## Introduction

The healing process of skin wounds requires precise regulation of inflammation, angiogenesis, cell migration, and proliferation, as well as extracellular matrix reorganization. Under normal circumstances, these processes are well-coordinated, taking place and resolving promptly to restore skin homeostasis (1, 2). However, excessive mechanical stretching at the site of a skin wound may interfere with the course of wound healing, leading to the formation of enlarged scars (3), and hypertrophic scars or keloids in predisposed individuals (4–6). Keloids, typically triggered after skin injury, are bulging scar tissues that extend beyond the boundary of the original wound, composed mainly of hyper-proliferative fibroblasts, excessive collagen deposited by fibroblasts, and neo-vasculature. Body areas subject to heightened mechanical stretching, such as the anterior chest and shoulders, are prone to keloid formation (7). Moreover, the local distribution of tension gradient dictates the direction of keloid progression (8, 9), indicating a prominent effect of mechanical stimulation on keloid pathogenesis. However, the identity of the major mechano-responsive cells in keloid tissues and their cellular characteristics underlying the disease progression remain incompletely understood.

Wound healing studies in rat and mouse models suggest that fibroblasts are mechano-perceptive (10–13). Using a rat tail stretching model (14), it has been shown that mechanical stretching up-regulates and activates fibroblasts’ PIEZO1, a mechano-sensitive calcium channel that relays mechanical stimuli by producing calcium signals, resulting in enlarged scar area and aggravated collagen deposition (11). Utilizing a full-thickness mouse dorsal wound model combined with a screw-driven stretching apparatus (15), the critical role of Engrailed 1 (EN1)-positive fibroblasts in the formation of mechanically induced enlarged scars has been pinpointed. Upon mechanical stretching, the EN1-positive fibroblasts activate FAK, a non-receptor tyrosine kinase that associates with integrins, transducing extracellular mechanical cues to the Rho/Rho-associated protein kinase (ROCK) signaling and the downstream transcription factor YAP (12). Pharmacological inhibition of PIEZO1 or YAP/TAZ with GsMTx4 (11) or verteporfin (12) in stretched rodent wound healing models reduced scarring. Although rodent models cannot fully recapitulate the development of keloids, pharmacological inhibition or knockdown of YAP/TAZ or PIEZO1 in keloid fibroblasts attenuated a series of pro-fibrotic phenotypes *in vitro*, including fibroblasts’ proliferation, contraction, migration, and collagen synthesis. Moreover, elevated protein levels of YAP/TAZ and PIEZO1 have been observed in clinical specimens of keloids compared to normal human skin (16, 17). Together, these results suggest a contribution of mechano-transduction in fibroblasts to keloid pathogenesis.

The application of single-cell RNA sequencing (scRNA-seq) technologies to human keloid specimens revealed four major fibroblast subtypes, namely mesenchymal, inflammatory, papillary, and reticular (18–23). Among these subtypes, the mesenchymal fibroblasts are amplified in keloid lesions and able to promote collagen production of non-mesenchymal fibroblasts via POSTN paracrine signaling (18). Spatial transcriptomic (ST) analysis has shown that some mesenchymal fibroblasts localize near vascular endothelial cells to facilitate mesenchymal activation of the vascular endothelium, promoting keloid angiogenesis (21). However, whether there is a mechano-responsive fibroblast group in human keloid tissues, its relationship with the existing four major fibroblast subtypes, and its cellular plasticity upon mechanical stretching, remain unexplored.

In this study, by coupling scRNA-seq data of keloid specimens with RNA sequencing (RNA-seq) data of stretched keloid fibroblasts, we unveil that CD74^+^ fibroblasts serve as the pivot of stretch-induced keloid progression, driven by their proliferative and pro-angiogenic capacities. Mechanical stretching stimulated the proliferation of CD74^+^ fibroblasts through PIEZO1-mediated calcium influx and subsequent activation of ERK and AKT pathways, likely contributing to their expansion in the leading edge of keloids. The CD74^+^ fibroblasts directly interacted with vasculature in keloid specimens, exerting pro-angiogenic functions through the secretion of a variety of pro-angiogenic cues. Together, we report the critical role of CD74^+^ fibroblasts in keloid pathogenesis, opening up new opportunities for the treatment of this non-lethal but distressing skin disorder of prevalence.

## Results

### Fibroblasts in Keloid are Mechano-responsive

To identify cell types responsive to mechanical stretching in keloid, analysis on an integrated dataset was conducted. Fastq files from five publicly available datasets comprising 13 keloid, 7 normal scar, and 10 normal skin samples were downloaded from respective repositories (Supplementary table 1), processed using scanpy (24, 25), and integrated with the BBKNN algorithm (26) to generate a dataset comprising 210,767 cells. Following unbiased Leiden clustering (27), 21 clusters were obtained (Figure 1A) and annotated into 12 cell types based on their respective marker genes (Figure 1B). Fibroblasts, endothelial cells, and keratinocytes emerged as the three major cell types in this cell atlas, constituting 34.5%, 20.6%, and 19.9% of all cells, respectively. Comparison of the relative abundance of cell types among sample groups revealed the expansion of both endothelial cells and fibroblasts in keloid (Figure 1C). Endothelial cells, being the most prominent cell types expanded, comprised 26% of total cells in keloid samples, compared to 17% of that in normal scar and 12% in normal skin, respectively. Fibroblasts also showed considerable expansion, comprising 38% of total keloid cells and 37% of total cells in normal scar, compared to 25% of that in normal skin (Figure 1C and Supplementary table 2). This finding corroborated previous reports of enhanced angiogenesis and hyper-proliferated fibroblasts in keloid. To evaluate the responsiveness to mechanical stretching of all cell types, a curated gene set (GO: Response to mechanical stimulus, GO:0009612) was used (28) to calculate a mechanical score for each cell type. As anticipated, the mechanical score of fibroblasts was significantly higher than that of any other cell type (Figure 1D). Moreover, within the fibroblasts, the score of the keloid population was higher than that of scar or skin (Figure 1E), indicating that the fibroblasts are exposed to an environment with increased mechanical stretching in keloid than in normal scar or skin.

**Figure 1.**
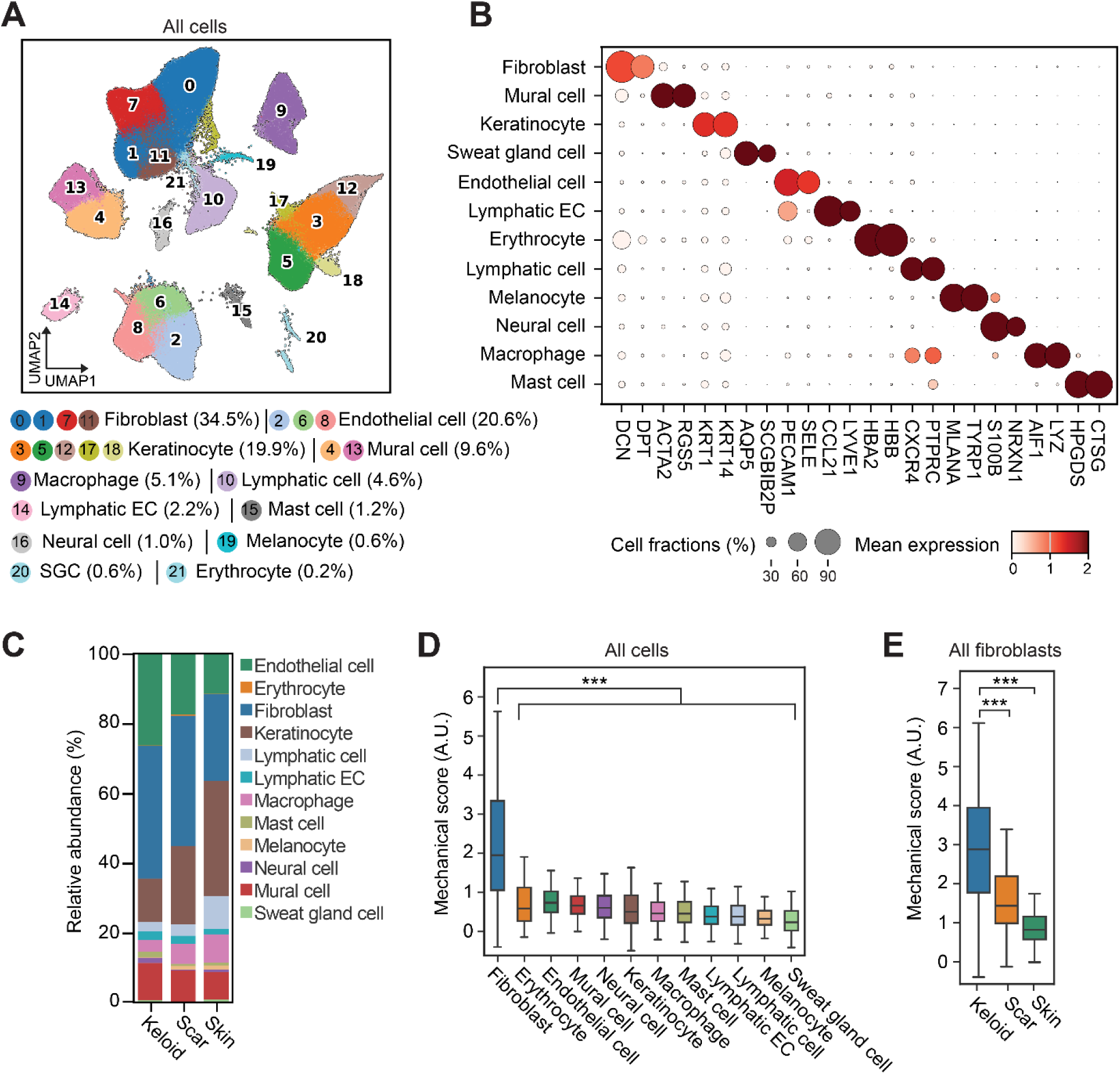
Fibroblasts are mechano-responsive in keloid. (A) UMAP projection and unbiased clustering of all cells included in the analysis with their relative abundance. (B) Dot plot shows the marker genes of each cluster. (C) Stacked bar plot shows the relative abundance of cell clusters in keloid, scar, and skin. (D) Boxplot shows the mechanical score of each cell cluster. (E) Boxplot shows the mechanical score of fibroblasts in keloid, scar, and skin. *** Denotes *P* ≤ 0.001, Student’s t test. A.U. denotes arbitrary unit.

### Identification of CD74^+^ Fibroblasts as a Candidate Mechano-responsive Fibroblasts Subtype in Keloid

To further dissect the heterogeneity of the fibroblast population, all fibroblasts (a total of 72,767 cells) were re-clustered to obtain 11 fibroblast subtypes with different marker genes (Figures 2A-2B). Since keloid fibroblasts were previously divided into four categories with distinct functions (18–23), we examined the relationship between our fibroblast subtypes and the prior classification. Subtypes 0, 1, 2, and 3 in our study were positive for POSTN expression, a marker of the previously defined mesenchymal fibroblasts (Supplementary figure 1A). Four other subtypes identified in our analysis (subtypes 4, 8, 9, and 10) could be categorized into secretory-reticular (subtype 4), secretory-papillary (subtype 8), and inflammatory fibroblasts (subtypes 9 and 10), according to the expression of their representative marker genes (Supplementary figure 1A). However, subtypes 5, 6, and 7, which collectively accounted for 5.7% of all fibroblasts, did not fit into these four categories (Figure 2A and Supplementary figure 1A). Among the newly identified 11 fibroblast subtypes, only subtypes 0, 1, 2, and 3 showed a proportional expansion in keloid samples (Figure 2C and Supplementary table 3), implying their potential pathological significance.

**Figure 2.**
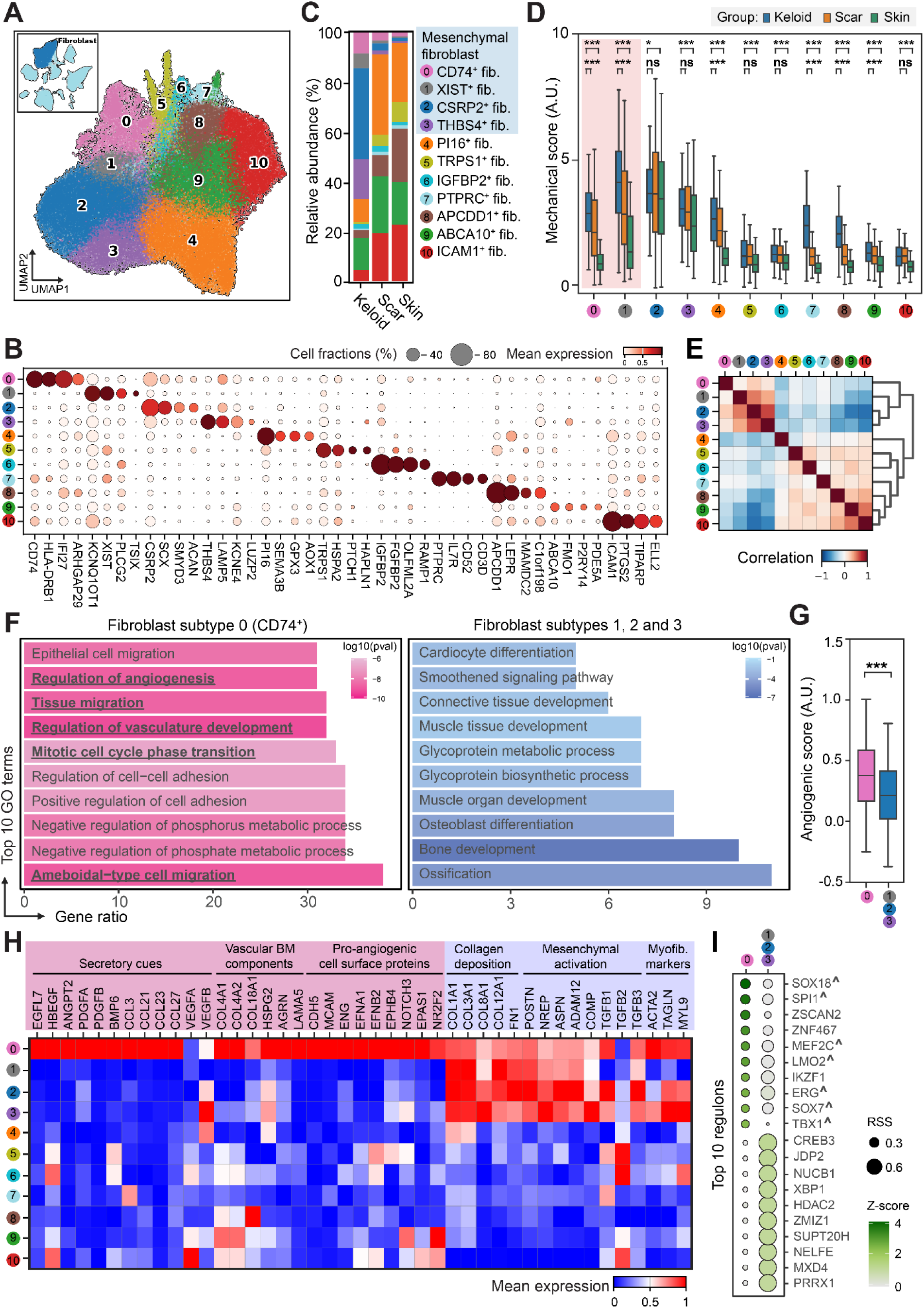
CD74^+^ fibroblasts respond to mechanical stretching and exhibit pro-angiogenic capacity. (A) UMAP projection and unbiased clustering of all fibroblasts. Top-left inset denotes the location of fibroblast cluster in Figure 1A. (B) Dot plot shows the marker genes of each fibroblast subtype. (C) Stacked bar plot shows the relative abundance and selected marker gene of each fibroblast subtype in keloid, scar, and skin. (D) Boxplot shows the mechanical score of each fibroblast subtype in keloid, scar, and skin. * Denotes *P* ≤ 0.05, *** denotes *P* ≤ 0.001, ns denotes *P* > 0.05, Student’s t test. (E) Correlation matrix and dendrogram of all fibroblast subtypes. (F) GO enrichment plots for genes up-regulated in fibroblast subtype 0 (left) and genes up-regulated in subtypes 1,2 and 3 (right). (G) Boxplot shows the angiogenic score of fibroblast subtype 0 versus subtypes 1,2 and 3 in keloid, scar, and skin. *** Denotes *P* ≤ 0.001, Student’s t test. (H) Heatmap shows the expression of selected pro-angiogenic genes in all fibroblast subtypes. (I) Dot plot shows the top 10 regulons predicted by pySCENIC in fibroblast subtype 0 versus subtypes 1,2 and 3 in keloid, scar, and skin. ∧ denotes pro-angiogenic transcription factors. A.U. denotes arbitrary unit.

Previous studies in mice have suggested *EN1*, *PTK2* (encoding FAK), and *JUN*, might serve as markers for mechano-responsive fibroblasts during wound healing (10, 12, 29). However, the expression of these genes failed to specifically highlight any of the fibroblast subtypes (Supplementary figure 1B). To directly assess the mechanical responsiveness, the mechanical score of all fibroblast subtypes was calculated. A significantly higher mechanical score was collectively observed in subtypes 0, 1, 2, and 3 (Supplementary figure 1C), with the scores of subtypes 0 and 1 exhibiting a marked increase in keloid compared to both scar and skin (Figure 2D), concentrating our research attention to these two subtypes. Upon examining the cell abundances, we noted that the number of cells belonging to subtype 0 was greater than that of subtype 1, comprising 8.9% in keloid and 6.5% in all fibroblasts. In comparison, fibroblast subtype 1 accounted for 5.9% of fibroblasts in keloid and 3.8% in all fibroblasts. The expression of multiple proliferation markers, including *MKI67*, *TOP2A*, *CENPE*, *CENPF*, and *MCM* genes (30), was higher in subtype 0 than that of other fibroblast subtypes (Supplementary figure 1D), indicating that subtype 0 was undergoing active proliferation. Additionally, correlation analysis of all fibroblast subtypes showed that subtypes 1, 2, and 3 were transcriptionally similar to each other, while subtype 0 was relatively distinct (Figure 2E). Given that fibroblast subtype 0 was abundant, expanded in keloid, and manifested elevated mechanical score, we termed this cell population CD74^+^ fibroblasts using one of its representative marker genes encoding a cell surface membrane protein and conducted further functional examination.

### CD74^+^ Fibroblasts, likely Originated from Myeloid Cells, Harbor Strong Pro-angiogenic Capacity

To unveil the unique functionality of CD74^+^ fibroblasts, a differential gene expression analysis of subtype 0 versus subtypes 1, 2, and 3 from all samples was conducted, resulting in 501 up-regulated and 116 down-regulated genes in subtype 0, respectively (Supplementary table 4). These genes were subjected to further functional enrichment analysis, revealing biological processes relevant to angiogenesis, cell migration, and cell proliferation among the top 10 enriched terms, highlighting the pro-angiogenic and proliferative capacities of the subtype 0 fibroblasts (Figure 2F). Consistently, the pro-angiogenic capacity of subtype 0 fibroblasts, estimated by scoring with an angiogenic gene set (GO: Angiogenesis, GO:0001525), was significantly higher than that of subtypes 1, 2, and 3 (Figure 2G). Examination of these 501 up-regulated genes in CD74^+^ fibroblasts revealed the involvement of several possible pro-angiogenic mechanisms, including secretion of various pro-angiogenic cues, production of vascular basement membrane components, and expression of pro-angiogenic signaling proteins on the cell surface (Figure 2H). These results indicated that CD74^+^ fibroblasts may contribute to keloid angiogenesis in a multi-factorial manner comprising fibroblast-to-endothelium paracrine signaling, stabilization of immature vessel structures, and direct support based on cell-cell contact, which is analogous to the pro-angiogenic strategy adopted by the vascular cancer-associated fibroblast (vCAF) in the tumor microenvironment (31, 32). Previous studies, including scRNA-seq and analysis on fibroblasts extracted from keloid lesions, have ascribed the pro-angiogenic effects exerted by keloid fibroblasts to enhanced VEGF, TGF-β, and ephrin signaling towards endothelial cells in keloid (18, 20, 33). Unexpectedly, among these three previously considered culprits in keloid angiogenesis, only ephrin ligands (*EFNA1* and *EFNB2*) exhibited a specific expression in CD74^+^ fibroblasts, while TGF-β ligands (*TGFB1*, *TGFB2*, and *TGFB3*) showed an indistinctive expression between CD74^+^ fibroblasts and the rest of the mesenchymal fibroblasts. As for VEGF signaling, whereas *VEGFB* was highly expressed in fibroblast subtype 3, *VEGFA* was highly expressed in fibroblast subtype 10, which were inflammatory fibroblasts whose relative cell abundance was negligible in keloids. This observation is in accordance with the spatial transcriptomic (ST) data of keloid specimens, where the ST spots positive for VEGF receptor-ligand interactions were scarce (21). Apart from the above-mentioned pro-angiogenic signals, *ANGPT2*, *PDGFA*, *PDGFB*, and multiple C-C motif chemokine ligands (*CCL3*, *CCL21*, *CCL23*, and *CCL27*) were found to be specifically expressed in CD74^+^ fibroblasts, representing a previously undiscovered route of angiogenesis in keloid. Again, this delineation emphasized the potential pro-angiogenic role of CD74^+^ fibroblasts in keloid. Regarding other pathogenic factors besides angiogenesis, such as collagen deposition, production of mesenchymal signals, and signs of myofibroblast activation, CD74^+^ fibroblasts exhibited little difference from the rest of the mesenchymal fibroblasts (Figure 2H).

Since CD74^+^ fibroblasts were distinguished from the rest of the mesenchymal fibroblasts in their pro-angiogenic and proliferative capacity, we sought to probe into their underlying mechanism. We conducted a regulon analysis (34) to explore the transcription factors that were specifically active in CD74^+^ fibroblasts. Expectedly, among the top ten specific transcription factors, seven of which governed pro-angiogenic programs (Figure 2I) as previously reported (35–41). For example, SOX18 can activate the expression of pro-angiogenic genes including *EFNB2* and *MMP7* (42), and pharmacological inhibition of SOX18 resulted in reduced vascular development in zebrafish larvae and tumor angiogenesis in murine models (43).

Surprisingly, we also noted the presence of several transcription factors crucial to myeloid differentiation, including *SPI1*, *MEF2C*, *LMO2*, *IKZF1*, and *ERG* (44–48). We hence sought to search for the origin of CD74^+^ fibroblasts. Trajectory inference analysis within fibroblast subtypes failed to identify differentiation processes related to the CD74^+^ fibroblasts. Since a variety of other cell types could give rise to fibroblasts (49), we computed a correlation matrix including all fibroblast subtypes as well as the rest of the cell clusters obtained from our first-pass clustering. On the matrix, we noted that the cell type showing the closest association with CD74^+^ fibroblasts was the macrophage (Supplementary figure 2A). This observation, together with the result of the regulon analysis, supports the possibility that macrophage-to-myofibroblast transition (MMT) may contribute to the appearance of CD74^+^ fibroblasts in keloid. The occurrence of MMT in keloid has not been documented. However, in a murine skin wound healing model (50), a unilateral ureteral obstruction (UUO)-induced murine renal fibrosis model (51), and a myocardial ischemia=reperfusion (MIR)-induced murine heart fibrosis model (52) where MMT activities were discovered, fibroblasts derived from MMT were characterized by *Lyz2*, *Cd206/Mrc1*, and *S100A9* expression, respectively. In CD74^+^ fibroblasts, specific expression of these three MMT marker genes, as well as a variety of lineage-specific markers of macrophages, was well preserved (Supplementary figure 2B), reemphasizing the possibility that CD74^+^ fibroblasts in keloid are originated from myeloid cells through MMT.

### Enrichment of CD74^+^ Fibroblasts in the Keloid Leading Edge and Their Interaction with Vasculature

Generally, the leading edge of keloids, which actively progresses and invades into adjacent normal skin, experiences more mechanical stretching than the quiescent center. Accordingly, we hypothesized that CD74^+^ fibroblasts were more abundantly distributed in keloid edges than in keloid centers as a response to the mechanical microenvironment. To validate this hypothesis, surgical keloid specimens were obtained from three patients, and the specimens were segmented into the leading edge and the center based on their direction of growth (Figure 3A). Immunofluorescence staining was then conducted on cryosections of keloid specimens. Initially, using CD31 as a marker for vasculature, we compared the vasculature area between the leading edge and the center of keloids. The average vasculature area was found to be 10% in the leading edge and 3% in the keloid center, confirming the previously reported (53) higher vasculature density at the leading edge of keloids compared to the center (Figure 3B). Subsequently, employing PDGFRA as a pan-fibroblast marker (18), we compared the proportion of CD74^+^ fibroblasts, identified as cells double-positive for PDGFRA and CD74 staining, between the leading edge and the center of keloids. As anticipated, at the leading edge of keloids, CD74^+^ fibroblasts accounted for an average of 20% of all fibroblasts, significantly higher than the 5% observed in the keloid center (Figures 3C-3D). Moreover, interactions between CD74^+^ fibroblasts and vasculature were observed (Figures 3C and 3E), indicating a potential role for CD74^+^ fibroblasts in facilitating angiogenesis through direct interaction with vasculature, echoing the findings from our single-cell analysis.

**Figure 3.**
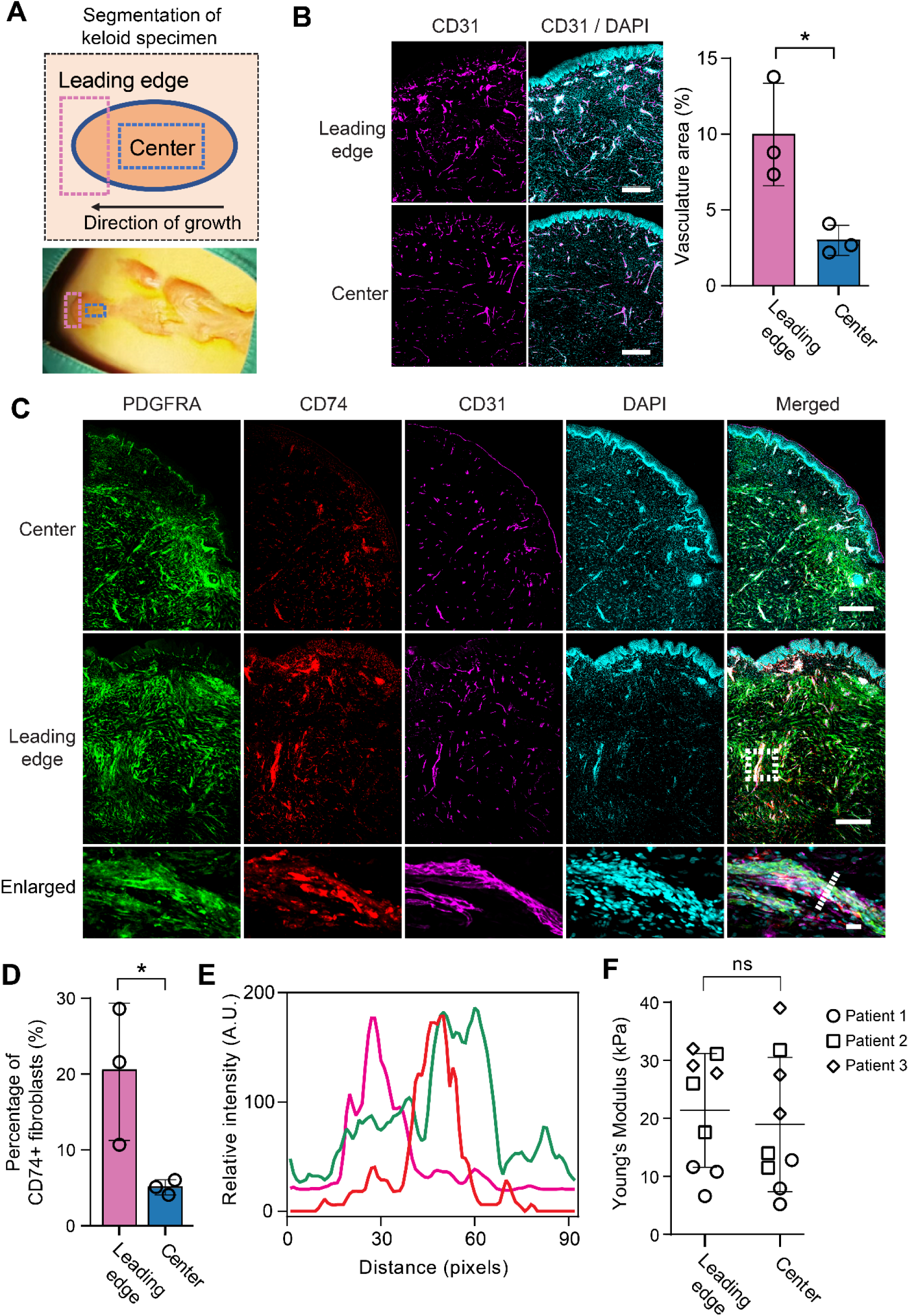
Enrichment of CD74^+^ fibroblasts in the keloid edge and interaction with vasculature. (A) Schematic of keloid segmentation strategy and gross photograph of a keloid specimen overlaid with segmentations of the leading edge and the center (bottom). (B) Representative immunofluorescence images (left) and quantification (right) of keloid cryosections stained with anti-CD31 antibody and DAPI. * Denotes *P* ≤ 0.05, Student’s t test. Scale bar: 500 µm. (C) Representative immunofluorescence images of keloid cryosections stained with anti-CD74 antibody, anti-PDGFRA antibody, anti-CD31 antibody, and DAPI. Dotted rectangle denotes an enlarged area. Dotted line denotes the line measured in figure 3E. Scale bar for leading edge and center: 500 µm. Scale bar for enlarged area: 20 µm. (D) Quantification of CD74^+^ fibroblast percentages of all fibroblasts in keloid leading edges and centers. * Denotes *P* ≤ 0.05, Student’s t test. (E) Intensity line scans of anti-CD74 antibody, anti-PDGFRA antibody, and anti-CD31 antibody immunostainings. (F) Measurement of Young’s modulus for three independent keloid specimens with atomic force microscopy. Statistical test was performed both collectively and patient-wise, and both showed no statistical significance. ns denotes P > 0.05, Student’s t test. A.U. denotes arbitrary unit.

Tissue stiffness can be perceived by residing cells, guiding the direction of cell migration in a phenomenon called durotaxis (54). To rule out whether the observed differences in CD74^+^ fibroblast distribution were influenced by alterations in tissue stiffness between the leading edge and the center of keloids, the stiffness of the leading edge and the center of keloids was measured using atomic force microscopy. Statistical tests, both collectively and patient-wise, yielded no significant differences (Figure 3F). This result suggested that tissue stiffness is not involved in the eccentric distribution of CD74^+^ fibroblasts in keloid tissues.

### Stretching Induces Proliferation of CD74^+^ Fibroblasts in Vitro

Given the observed proliferative capacity of CD74^+^ fibroblasts in single-cell data (Supplementary figure 1D), we hypothesized that mechanical stretching might enhance the proliferation of CD74^+^ fibroblasts in keloids, contributing to the expansion of CD74^+^ fibroblasts in keloid samples. To investigate this hypothesis, we utilized two fibroblast cell lines, keloid fibroblasts (KF) and human foreskin fibroblasts (HFF). Initially, we assessed the presence of CD74^+^ fibroblasts in these two cell lines using flow cytometry. CD74^+^ fibroblasts constituted 2.15% of all cells in KF but were scarcely present in HFF, accounting for 0.59% of all cells (Figure 4A), reflecting the expansion of CD74^+^ fibroblasts observed in keloid samples. Additionally, we examined the transcriptomic profiles of HFF and KF by RNA-seq to validate the representativeness of these two cell lines to the fibroblast subtypes identified in single-cell data. RNA-seq revealed that KF up-regulated genes associated with fibroblast subtypes enriched in keloid samples, while HFF up-regulated genes associated with fibroblast subtypes enriched in normal scar or normal skin samples. Specifically, among 6410 genes that were up-regulated in KF, marker genes of fibroblast subtypes 0, 2, 3, along with pan-markers for mesenchymal fibroblasts (namely, *POSTN* and *NREP*), were found, corresponding to the fibroblast subtypes that were enriched in keloid samples. Among 6695 genes that were up-regulated in HFF, marker genes of fibroblast subtypes 1, 4, 5, 6, 7, 8, 9, and 10, were found, corresponding to the fibroblast subtypes that were mainly enriched in normal skin samples (Figure 4B). This result suggested that the KF and HFF partially recapitulated fibroblast subtypes in keloid and normal skin samples, respectively.

**Figure 4.**
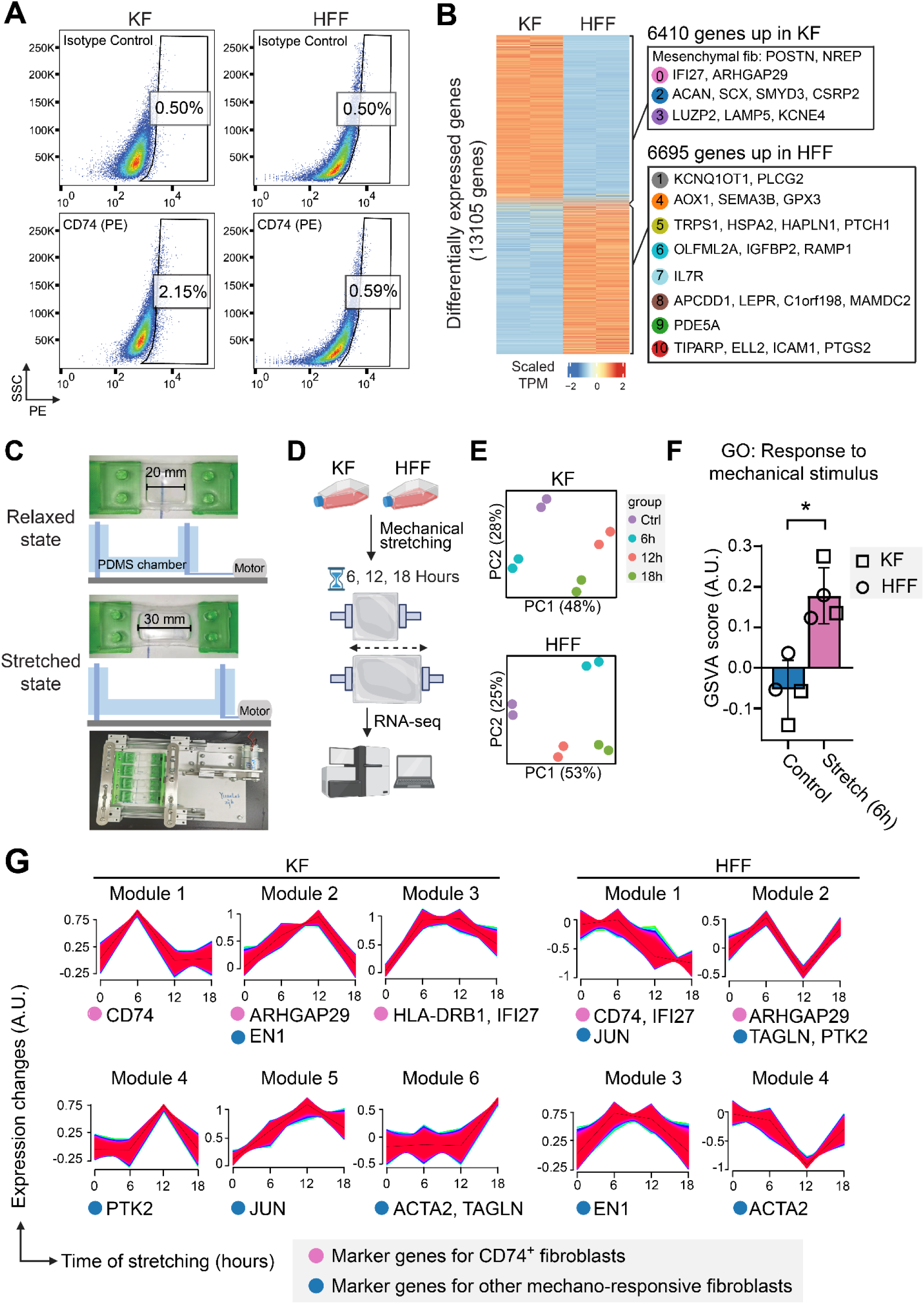
Mechanical stretching promotes the proliferation of keloidal CD74^+^ fibroblast. (A) Representative FACS plots for CD74^+^ cells in KF and HFF. (B) Heatmap shows the scaled TPM for differentially expressed genes in KF and HFF. The right panel shows all marker genes for fibroblast subtypes (figure 2B) that are included in differentially expressed genes. (C) Photograph and schematic of the stretching device used in cell stretching experiments. (D) Schematic depicting the procedure for cell stretching and sequencing. (E) PCA plots of all sequenced samples. (F) GSVA scores of KF and HFF cells stretched for 6 hours. * Denotes *P* ≤ 0.05, Student’s t test. (G) Time-series gene modules extracted from sequencing data by the Mfuzz algorithm. The presence of CD74^+^ fibroblast marker genes, myofibroblast markers, and other selected markers for mechano-responsive fibroblasts were annotated under respective gene modules. A.U. denotes arbitrary unit.

To investigate whether stretching could induce CD74^+^ fibroblast expansion *in vitro*, we stretched the KF and HFF cells on a custom-made cell stretching device and sent them for RNA-seq to assess the proportional changes of CD74^+^ fibroblasts. The custom-made cell stretching device, inspired by a previous study (55), was based on a PDMS elastomer and a direct current (DC) motor (Figure 4C). KF and HFF were stretched at 0.1 Hz for 6, 12, and 18 hours before RNA-seq (Figure 4D). Mechanical stretching significantly altered the transcriptomes of KF and HFF (Figure 4E). After 6 hours of stretching, both cell lines exhibited significantly elevated mechanical scores as calculated by gene set variation analysis (GSVA), indicating a robust response to stretching stimuli (Figure 4F). Then, using the Mfuzz algorithm (56), we clustered genes with similar expression change patterns over time into gene modules (Figure 4G). A total of 15 gene modules were obtained from RNA-seq data of each cell line. Examining the presence of marker genes for CD74^+^ fibroblasts within these modules allowed us to infer proportional changes of CD74^+^ fibroblasts. In stretched KF, all four marker genes for the CD74^+^ fibroblast population, namely *CD74*, *IFI27*, *HLA-DRB1*, and *ARHGAP29*, appeared in gene modules with elevated expression levels, e.g., modules 1, 2, and 3. On the contrary, in HFF, *CD74*, *IFI27*, and *ARHGAP29* were found in modules with declined or fluctuating expression levels, e.g., modules 1 and 2. The expression change of *HLA-DRB1* was insignificant in HFF as determined by the algorithm, therefore it was not shown. This result is in accord with the expected stretch-induced expansion of CD74^+^ fibroblasts. In order to gain a more comprehensive view of stretch-induced effects on fibroblasts, we also examined several marker genes for previously defined mechano-responsive or pro-fibrotic fibroblasts, namely, *EN1*, *PTK2*, and *JUN*, as well as myofibroblast markers, *ACTA2* and *TAGLN*, within these gene modules. Similar to marker genes of CD74^+^ fibroblasts, these genes appeared in gene modules with elevated expression levels, e.g., modules 2, 4, 5, and 6, in stretched KF. On the contrary, in HFF, the majority of these five genes appeared in modules 1, 2, and 4, whose gene expression levels declined or fluctuated, except for *EN1*.

To determine whether the selective proliferation of CD74^+^ fibroblasts contributed to the stretch-induced up-regulation of CD74^+^ fibroblast marker genes, both CD74^+^ and CD74^-^ fibroblasts in KF were purified with fluorescence-activated cell sorting (FACS), stretched for 6 hours, and subjected to BrdU labeling and subsequent immunofluorescence staining. After FACS sorting, cell purity was confirmed via live-cell imaging (Supplementary figure 3). The result of BrdU labeling revealed a higher proliferative percentage in the stretched CD74^+^ KF, compared to the stretched CD74^-^ KF (Figures 5A-5B). Therefore, it was delineated that the stretch-induced CD74^+^ KF expansion was partially achieved by the selective proliferation of this cell population in the mechanical microenvironment.

**Figure 5.**
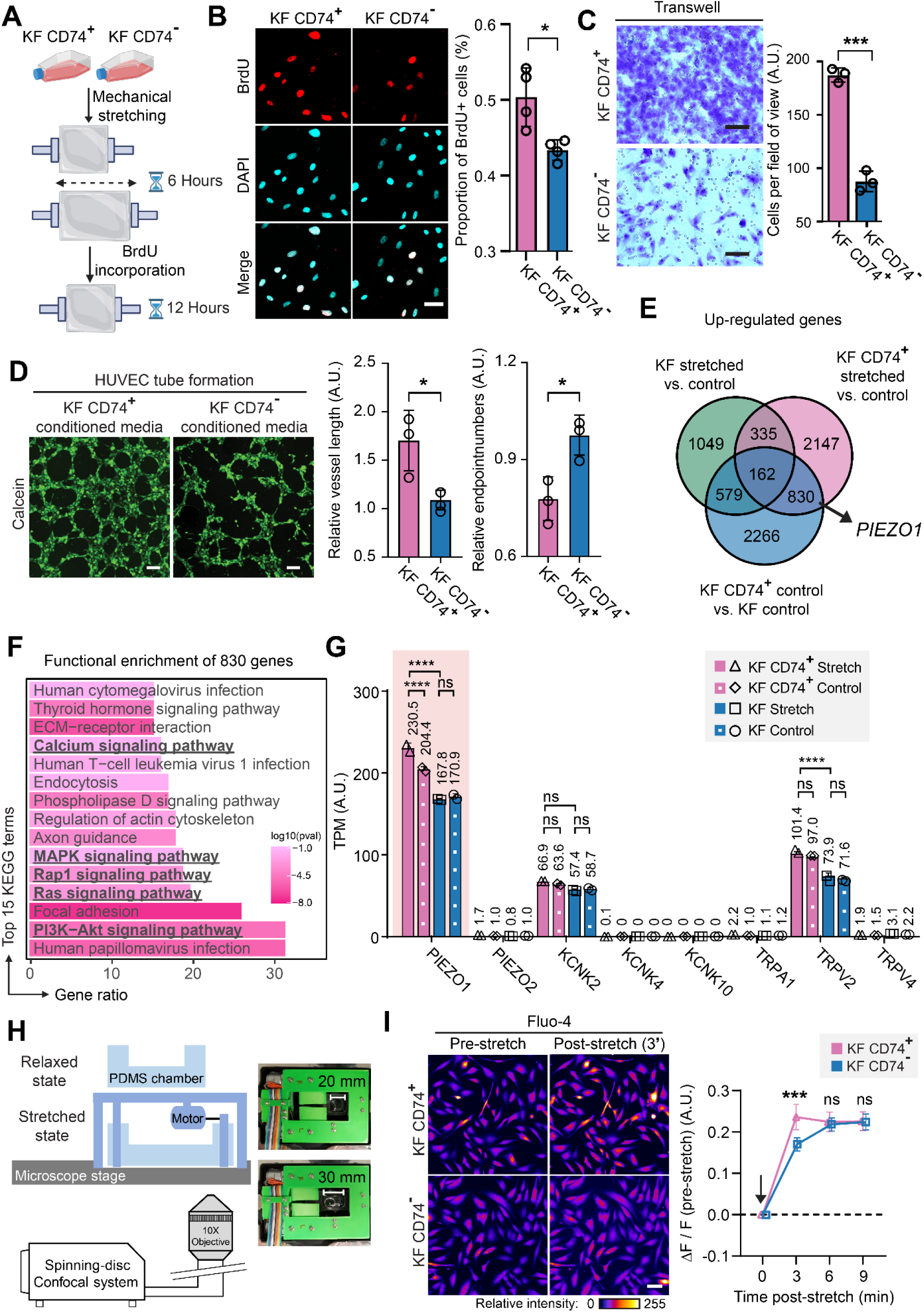
Stretching activates and up-regulates PIEZO1 in CD74^+^ fibroblasts. (A) Schematic depicting the experimental procedure for stretching and BrdU labeling of cells. (B) Representative images (left) of BrdU immunostaining and quantification (right) of BrdU^+^ cell proportions. * Denotes *P* ≤ 0.05, Student’s t test. Scale bar: 50 µm. (C) Representative images (left) and quantification (right) of transwell migration assay done with CD74^+^ and CD74^-^ KF. *** denotes *P* ≤ 0.001, Student’s t test. Scale bar: 100 µm. (D) Representative images (left) and quantification (right) of HUVEC tube formation assay done with conditioned culture media obtained from CD74^+^ and CD74^-^ KF. * Denotes *P* ≤ 0.05, Student’s t test. Scale bar: 200 µm. (E) Venn diagram shows the number of up-regulated genes in three sets of differential expression analysis. Arrow denotes the presence of PIEZO1 in a list of 830 genes that were up-regulated both in CD74^+^ KF (compared to KF) and in stretched CD74^+^ KF (compared to unstretched CD74^+^ KF) but not in stretched KF (compared to unstretched KF). (F) KEGG functional enrichment result of 830 genes. (G) Expression levels of all mechano-sensitive calcium channels in KF and CD74+ KF before and after stretching, as measured by TPM. **** Denotes *P* ≤ 0.0001, DESeq2 analysis. (H) Schematic (left) and photographs (right) of the stage-top cell stretching device used in the Fluo-4 experiment. (I) Representative Fluo-4 live cell fluorescence images (left) and quantification (right) of stretched CD74^+^ and CD74^-^ KF. For each cell, the Fluo-4 fluorescence signal (F) pre-stretch, and after 20 cycles of stretching for 3, 6, and 9 minutes, was recorded. Then, ΔF/F(pre-stretch) was calculated. Mean ± 95% confidence interval of the data was reported. *** Denotes *P* ≤ 0.001, ns denotes *P* > 0.05, Student’s t test. Arrow denotes the application of mechanical stretching. For CD74^+^ KF, n=224. For CD74^-^ KF, n=298. Scale bar: 50 µm. A.U. denotes arbitrary unit.

Our analysis of single-cell data emphasized the migratory and pro-angiogenic capacities of CD74^+^ fibroblasts in keloid. With purified CD74^+^ and CD74^-^ KF at hand, we were able to validate these two traits using transwell migration and tube formation assays, respectively. For transwell migration assay, CD74^+^ KF or CD74^-^ KF were added to the upper chamber of the transwell and incubated for 24 hours, then, the cells migrated through the membrane were quantified by staining with crystal violet. The result showed a greater migratory ability of CD74^+^ KF compared with CD74^-^ KF (Figure 5C). For tube formation assay, the conditioned media of CD74^+^ KF or CD74^-^ KF were added to the Human Umbilical Vein Endothelial Cells (HUVEC) on matrigel, and the results showed that CD74^+^ KF has greater pro-angiogenic capacity than CD74^-^ KF (Figure 5D). These phenotypic results aligned with our single-cell analysis.

### CD74^+^ Fibroblasts Sense and Respond to Mechanical Stretching through PIEZO1-Calcium-ERK/AKT Axis

To elucidate the molecular events responsible for stretch-induced CD74^+^ fibroblast proliferation, we conducted RNA-seq on mechanically stretched CD74^+^ KF. Then, we intersected the up-regulated genes from three sets of differential expression analyses: stretched KF vs. unstretched KF, stretched CD74^+^ KF vs. unstretched CD74^+^ KF, and unstretched CD74^+^ KF vs. unstretched KF. The intersection identified a list of 830 genes that were up-regulated both in CD74^+^ fibroblasts (compared to unsorted KF) and in stretched CD74^+^ fibroblasts (compared to unstretched CD74^+^ KF) but not in stretched KF (compared to unstretched KF) (Figure 5E). These genes were subject to functional enrichment analysis, which identified several terms related to the activation of ERK1/2, AKT as well as calcium signaling, implying the pivotal role of stretch-induced calcium channel activation in CD74^+^ fibroblast proliferation (Figure 5F). To identify the specific calcium channel that plays a leading role, a panel of known mechano-sensitive ion channels was examined in our data (57). The result showed that *PIEZO1*, *KCNK2*, and *TRPV4* were expressed in KF cells, while the rest manifested negligible expression levels. Among these three candidates, Both *PIEZO1* and *TRPV2* showed higher expression levels in CD74^+^ KF compared to unsorted KF, serving as mechano-sensors which could explain the sensitivity of CD74^+^ fibroblasts to stretching. Furthermore, the expression of *PIEZO1* was further elevated in stretched CD74^+^ KF, suggesting its role as a major mechano-sensor amplified by positive feedback mechanisms (Figure 5G). To visualize the stretch-induced calcium signals in CD74^+^ KF, we utilized a custom-made stretching device compatible with confocal microscopy (Figure 5H). CD74^+^ KF or CD74^-^ KF were seeded on the PDMS elastomer, preloaded with Fluo-4, an indicator of intracellular concentration of calcium ions, and stretched for 20 cycles. The elastomer was then mounted on the microscope stage and cells were imaged with confocal microscopy at the indicated timepoints. We observed a significantly stronger calcium signal in stretched CD74^+^ KF compared to CD74^-^ KF 3 minutes after stretching (Figure 5I).

Previous studies showed that in stretched pulmonary arterial endothelial cells, the activation of cellular ERK1/2 and AKT signaling pathways was downstream of PIEZO1 activation (58). Given that we also identified several terms relating to the activation of ERK1/2 and AKT pathways from the enrichment analysis (Figure 5F), we queried phosphorylation levels of ERK1/2 and AKT in stretched CD74^+^ KF and CD74^-^ KF cells with western blotting. For both cell types, the quantity of protein in stretched cells was normalized to that of unstretched cells to calculate the variation of a specific protein. The results showed a significantly higher increment of phosphorylated ERK1/2 and AKT in stretched CD74^+^ KF, compared to that in stretched CD74^-^ KF (Supplementary figure 4A), indicating the activation of ERK1/2 and AKT signaling downstream the PIEZO1-mediated calcium influx. To further pin down the role of PIEZO1 in stretch-induced CD74^+^ fibroblast proliferation, we knocked down PIEZO1 using small interfering RNA (siRNA) in CD74^+^ KF and CD74^-^ KF cells, followed by 6 hours of stretching and subsequent BrdU labeling and western blotting (Supplementary figures 4B-4C). Knockdown of about 40% of the PIEZO1 level was achieved in both CD74^+^ and CD74^-^ KF cells (Supplementary figure 4B). Upon PIEZO1 knockdown, stretch-induced increment of phosphorylation was weakened in CD74^+^ KF (Supplementary figure 4D). Moreover, stretch-induced proliferation of CD74^+^ KF was significantly diminished upon PIEZO1 knockdown (Supplementary figure 4E).

Combined, these data support a model for stretch-induced keloid progression, where mechanical stretching drives the expansion of pro-angiogenic CD74^+^ fibroblasts, mediated by PIEZO1 and downstream phosphorylation signals activated by calcium influx (Figure 6).

**Figure 6.**
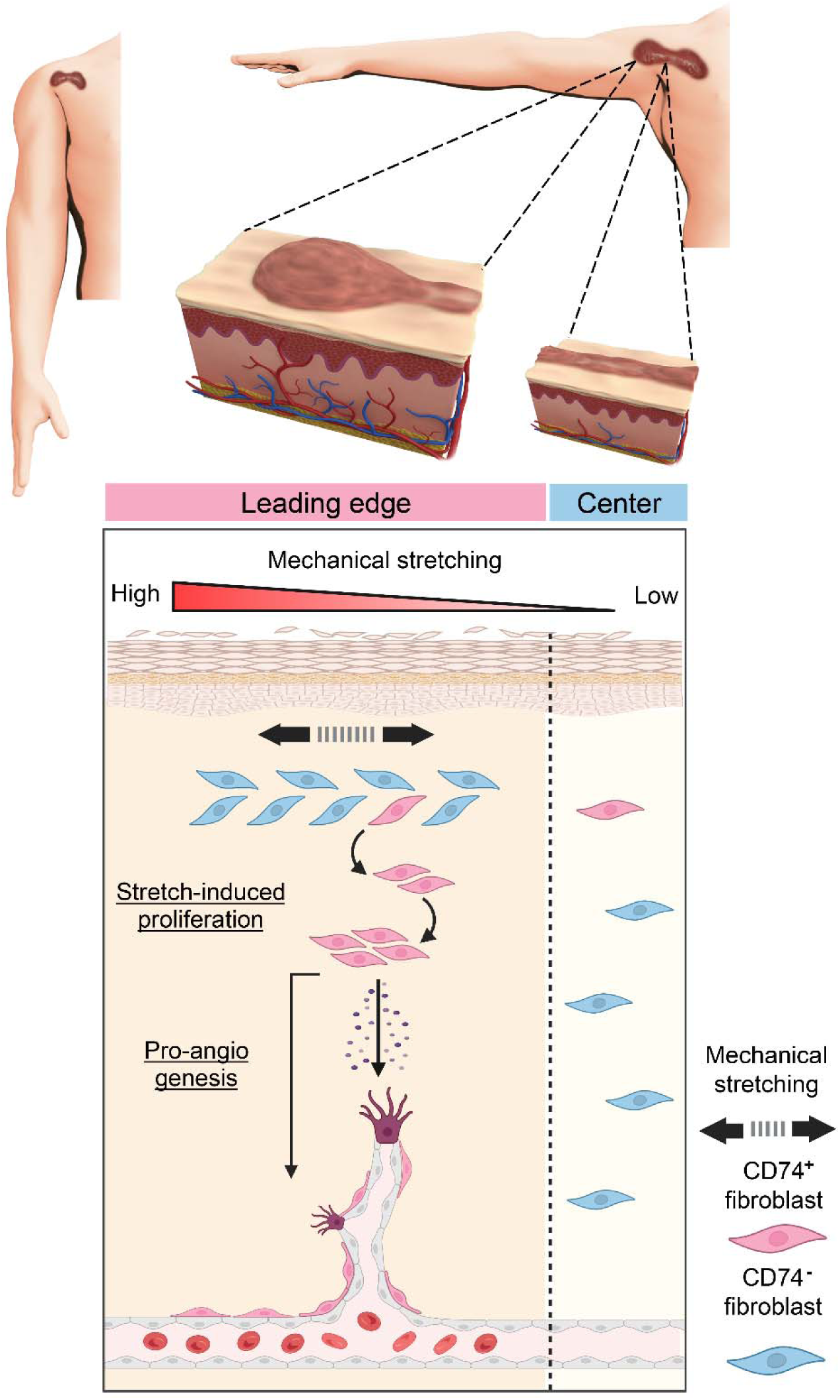
Working model of this study. Body parts subject to heightened mechanical stretching are prone to keloid formation, and the local distribution of stretching forces dictates the direction of keloid outgrowth, resulting in the actively progressing leading edge and the quiescent center. Stretch-induced keloid progression and neo-vascularization are two traits prominently manifested in the leading edge of keloids and facilitate keloid progression. Stretch-induced proliferation and pro-angiogenic capacity of CD74^+^ fibroblasts underlie these two traits. CD74^+^ fibroblasts exert pro-angiogenic activities through the secretion of pro-angiogenic cues and direct interaction with vasculature.

## Discussion

Keloids, bulging scar tissues invading into adjacent skin resulting from abnormal wound healing, can cause significant emotional and physical distress in patients (59). Keloids are preferentially formed in body areas subject to frequent mechanical stretching. There is an urgent need to pursue in-depth investigations into the mechanistic role of mechanical stretching on keloid pathogenesis in order to develop new therapies. In this study, we uncovered a distinct population of fibroblasts, the CD74^+^ fibroblasts, that manifest stretch-induced proliferation and pro-angiogenic capacity in keloid samples, establishing a relationship between mechanical stretching and neo-vascularization in keloid lesions. Our findings enhance the understanding of keloid pathogenesis and provide valuable insights into potential therapeutic interventions. Moreover, since the increased mechanical perturbation is a common feature found in many of the fibrotic diseases (60), it is worthy of investigation whether CD74^+^ fibroblasts also exist in other fibrotic diseases, promoting disease progression.

The fibro-proliferative and invasive nature of keloids necessitates a robust angiogenic mechanism at its command to transport oxygen and nutrients. Indeed, enhanced neo-vascularization activities of endothelial cells (53) as well as fibroblasts with stronger pro-angiogenic secretory phenotypes (33) were observed at the leading edge of keloids compared to keloid centers. The standard-of-care treatment modalities for keloids, including surgical excision, compression dressings, laser therapy, radiation, corticosteroids, and cytotoxic drugs, offer only limited efficacy with a high risk of recurrence (63). Many of these interventions are believed to exert therapeutic effects partly by inhibiting angiogenesis in keloid (41–43), however, their mode of action is unspecific to a large extent and incur considerable side effects (59), and the targeted inhibition of angiogenesis in keloid is yet to be accomplished. In this work, the pro-angiogenic capacity of CD74^+^ fibroblasts is unveiled, which may constitute a target for the future development of new anti-angiogenic therapy in keloid. Our analysis demonstrated that CD74^+^ fibroblasts specifically express ANGPT2, a member of the angiopoietin growth factor family whose up-regulation was reported in many human cancers (64). Since the mechanism of ANGPT2 in promoting angiogenesis is different from that of VEGFA (65, 66), dual inhibition of ANGPT2 and VEGFA has been proposed. Bispecific antibodies targeting ANGPT2/VEGFA, such as VA2 or Faricimab, approved by the FDA in 2022, have demonstrated satisfactory anti-angiogenic potency in cancer (67) as well as in retinal vascular diseases (68). Therefore, this may also represent a new avenue for keloid management.

Through analysis of single-cell data, we noted the possibility that macrophage to myofibroblast transition (MMT) may contribute to CD74^+^ fibroblasts in keloid. Since its discovery in renal fibrosis (51), evidence of MMT has been proposed in pulmonary fibrosis, cardiac fibrosis, and wound healing process (50–52, 61). However, the occurrence of MMT in keloid has not been documented. In this study, the specific expression of *LYZ2*, *CD206/MRC1*, and *S100A9*, which are markers for cells that underwent MMT, as well as a variety of lineage-specific markers of macrophages, has been found in CD74^+^ fibroblasts, supporting the possibility that CD74^+^ fibroblasts in keloid may, at least in part, originate from myeloid lineage. It’s remarkable that, in an investigation using a murine stretch-induced scarring model, local application of Fingolimod (FTY720), an immunosuppressant, decreased the number of macrophages and reduced the amount of micro-vessels in the wound bed, thereby reducing the gross scar area (62). Whether MMT gives rise to CD74^+^ fibroblasts in keloids, and whether mechanical stretching accelerates this process, are open questions, and further investigation is warranted.

Apart from keloids which are benign fibro-proliferative skin diseases, fibroblasts also serve as an essential component of the tumor microenvironment, influencing the initiation, progression, and treatment response of solid tumors (69). Intriguingly, we note that the phenotype of CD74^+^ fibroblasts that we identified in this study resembles a combination of vascular cancer-associated fibroblasts (vCAF) and antigen-presenting cancer-associated fibroblasts (apCAF), whose presence has been confirmed in multiple cancers (31, 32). Their resemblance includes (i) The marker genes of these cell types show a partial overlap, specifically, CD74^+^ fibroblasts express CD74, HLA-DRB1, MCAM, and NOTCH3, while apCAF expresses CD74 and HLA-DRB1, and vCAF expresses MCAM and NOTCH3; (ii) both CD74^+^ fibroblasts in keloid and vCAF in tumor microenvironment exhibit pro-angiogenic features. This phenomenon raises two questions. First, whether the specific expression of CD74 and HLA-DRB1 in CD74^+^ fibroblasts, which are key components of the major histocompatibility complex class II (MHC II) complex, indicates an antigen presentation capability of these cells is unknown. Second, considering the fact that the intra-tumoral environment where CAFs reside in is also brimming with mechanical stimulus (70), it is compelling to speculate whether vCAF undergoes proliferation upon mechanical stimulus, thereby contributing to neo-vascularization in tumors.

## Materials and Methods

All experimental protocols, manufacture of devices used in this study, and sequences of primers and oligonucleotides are provided in the SI Appendix.

### Statistical Analysis

Statistical analyses were performed using GraphPad Prism software (version 9.5.1, GraphPad) or R software (version 4.2.1). Student’s t tests and ANOVA were used to determine statistical differences between two conditions and among multiple conditions, respectively. Two-sided testing was performed for each analysis, and a *P* value less than 0.05 was considered statistically significant. Unless otherwise noted, the mean ± SD is reported.

## Data and Code Availability

All sequencing data generated in this study are publicly available through the Gene Expression Omnibus (GEO), under accession number GSE262112. All custom scripts and data included in this research are available from the authors on request.

## Acknowledgments

We thank Prof. Congying Wu and Dr. Xin Yi for their assistance with AFM measurements. We also thank Dr. Yantao Xu, Dr. Zixi Jiang, Dr. Zhenjun Deng, Dr. Qing Yu, and Dr. Dingyu Wu for their invaluable advice. This project has been supported by the National Natural Science Foundation of China (grants 32370821, 32170821, and 92153301 to K.Y), National Key Research and Development Program of China (2021YFC2701200), and Department of Science & Technology of Hunan Province (grants 2023RC1028 and 2023SK2091 to K.Y).

---

**Supplementary figure 1.**
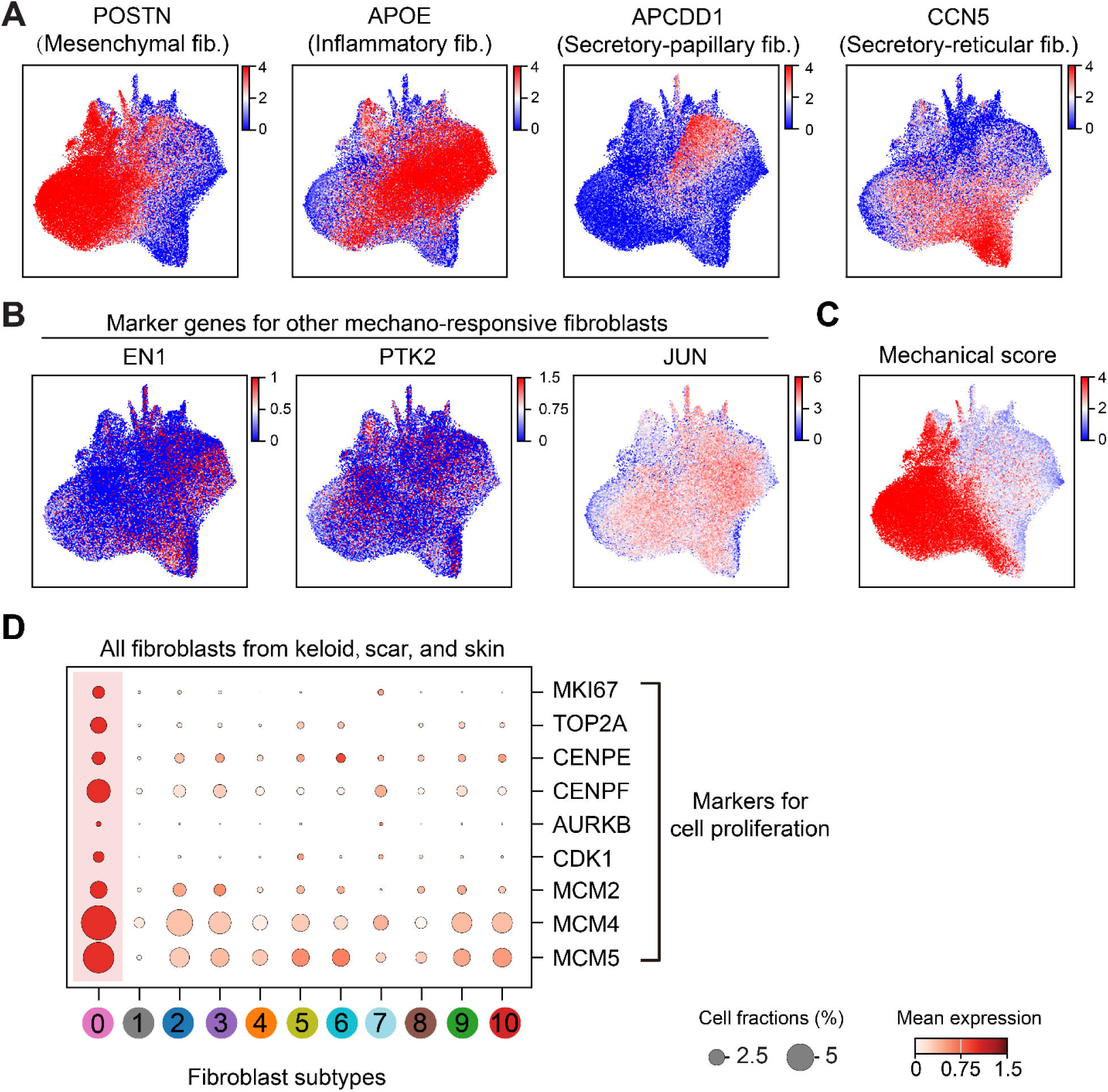
Relative expression levels of selected marker genes in all fibroblast subtypes from keloid, scar, and skin. (A) UMAP overlay of the relative expression level of fibroblast marker genes in all fibroblasts. (B) UMAP overlay of the relative expression level of mechano-responsive fibroblast marker genes in all fibroblasts. (C) UMAP overlay of mechanical scores in all fibroblasts. (D) Dot plot shows the expression level of cell proliferation markers in all fibroblast subtypes.

**Supplementary figure 2.**
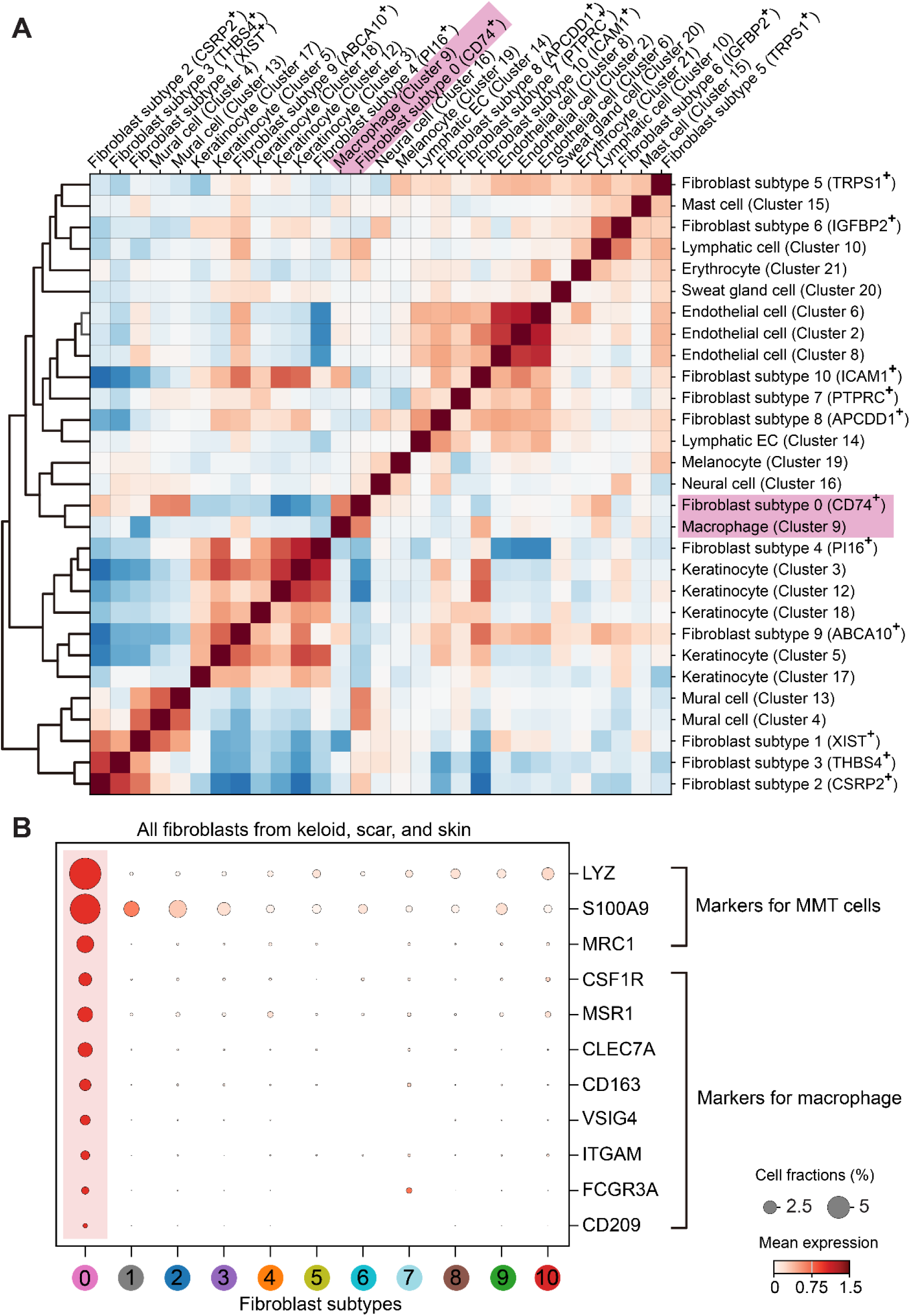
CD74^+^ fibroblasts may originate from myeloid-lineage cells. (A) Correlation matrix of all cell types derived from first-pass clustering and all fibroblast subtypes. (B) Dot plot shows the expression level of MMT and macrophage marker genes in all fibroblast subtypes.

**Supplementary figure 3.**
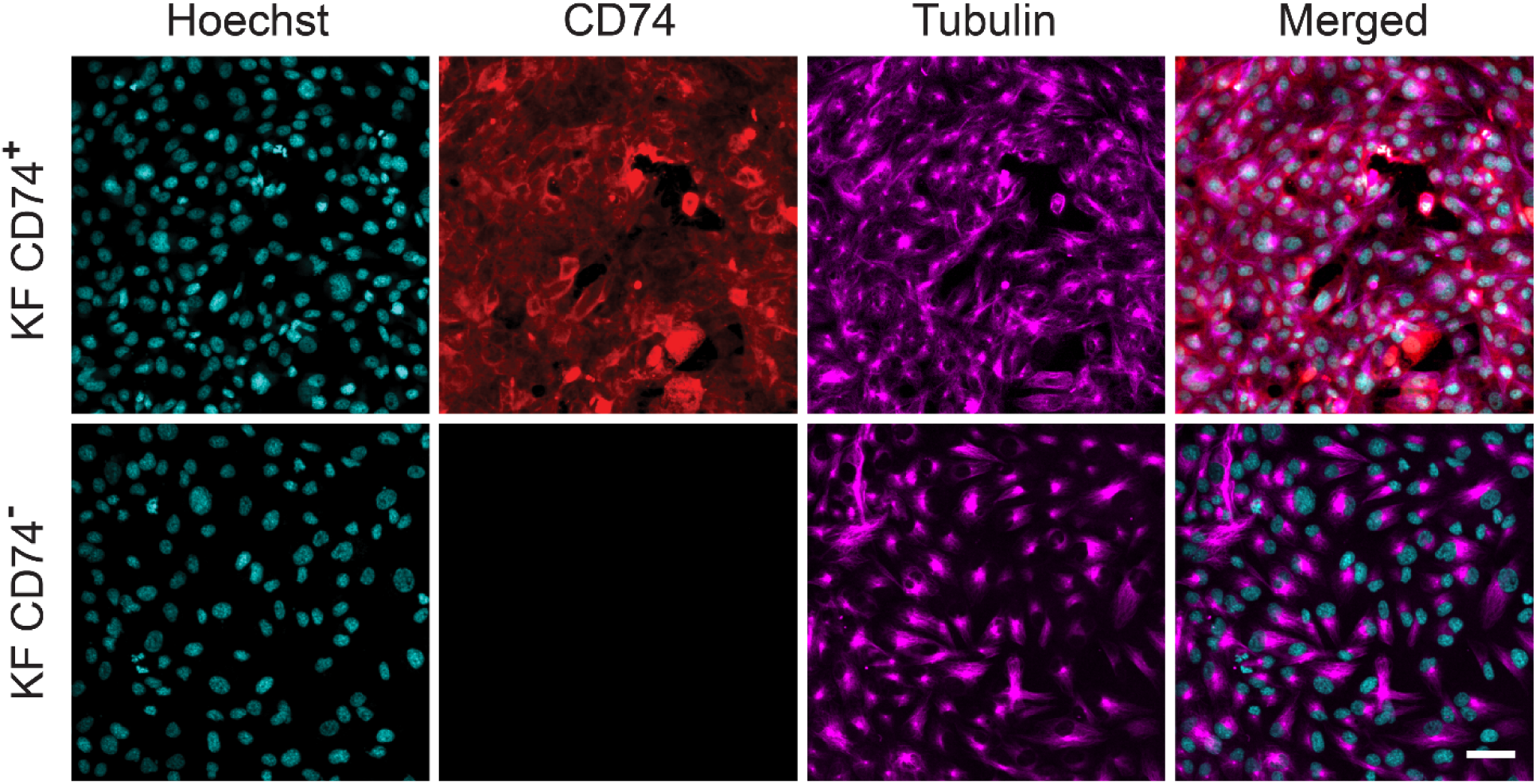
Live-cell imaging of FACS-purified CD74^+^ fibroblasts and CD74^-^ fibroblasts verified their purity. Cells were stained with anti-CD74 antibody, SPY650-tubulin, and Hoechst 33342. Scale bar: 50 µm.

**Supplementary figure 4.**
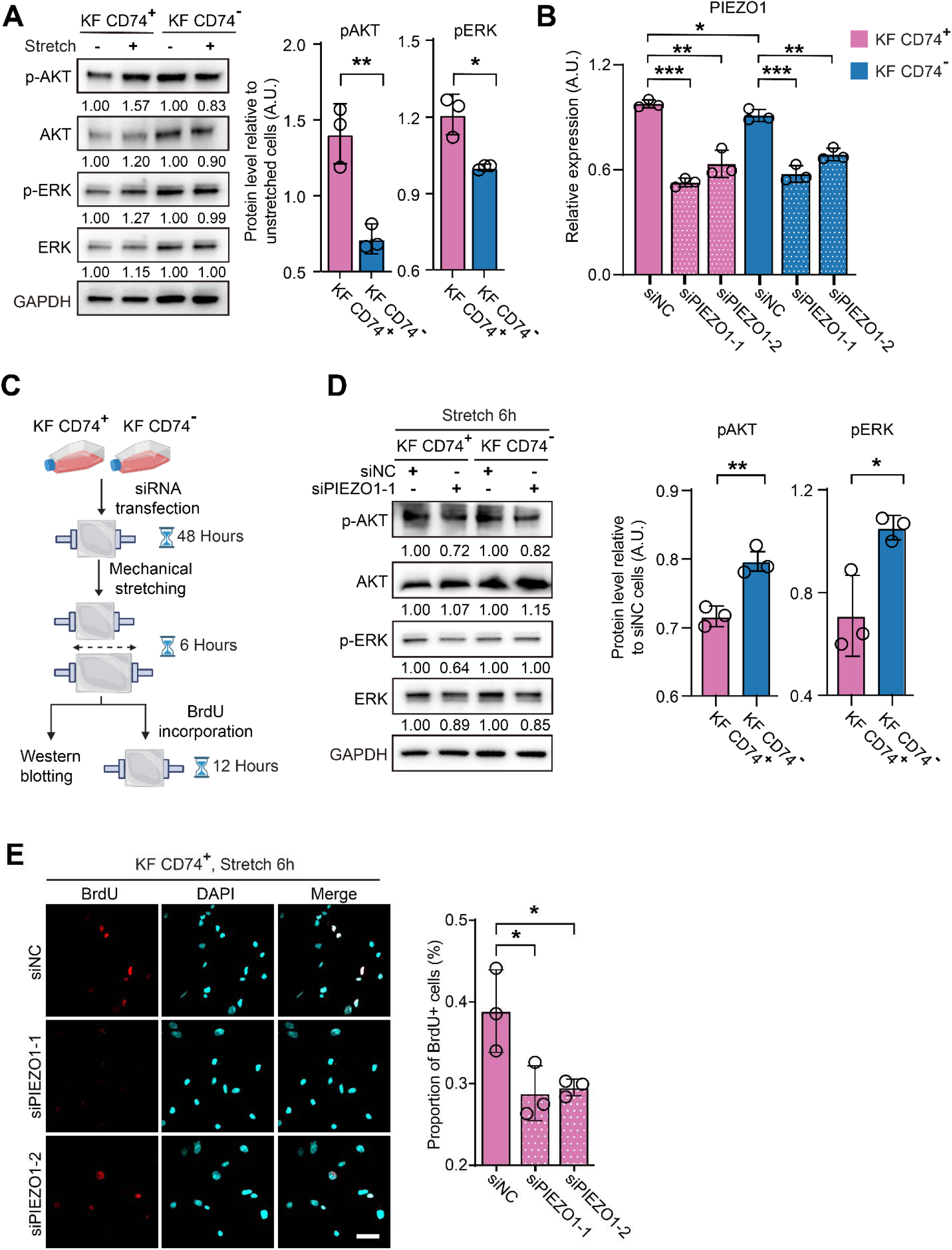
Activation of ERK and AKT in CD74^+^ fibroblasts upon mechanical stretching. (A) Representative western blot images (left) and quantification (right) of stretched and unstretched CD74^+^/CD74^-^ KF. The quantification was done independently on CD74^+^ and CD74^-^ KF as follows: for each target protein, the band intensity was firstly normalized with GAPDH, then the protein level in stretched cells relative to unstretched cells was calculated. * Denotes *P* ≤ 0.05, ** denotes *P* ≤ 0.01, Student’s t test. (B) Relative expression levels of PIEZO1 in cells transfected with siPIEZO1-1, siPIEZO1-2, or siNC, as measured by qRT-PCR. (C) Schematic depicting the experimental procedure of mechanical stretching of cells transfected with siPIEZO1 or siNC. (D) Representative western blot images (left) and quantification (right) of stretched CD74^+^/CD74^-^ KF transfected with siPIEZO1 or siNC. The quantification was done independently on CD74+ and CD74-KF as follows: for each target protein, the band intensity was firstly normalized with GAPDH, then the protein level in cells transfected with siPIEZO1-1 relative to cells transfected with siNC were calculated. * Denotes *P* ≤ 0.05, ** denotes *P* ≤ 0.01, Student’s t test. (E) Representative BrdU immunostainings (left) and quantification (right) of stretched CD74^+^/CD74^-^ KF transfected with siPIEZO1 or siNC. * Denotes *P* ≤ 0.05, Student’s t test. Scale bar: 50 µm.

## Notes

### Competing Interest Statement

The authors have declared no competing interest.

## References

1. G. C. Gurtner, S. Werner, Y. Barrandon, M. T. Longaker, Wound repair and regeneration. Nature 453, 314–321 (2008).

2. O. A. Peña, P. Martin, Cellular and molecular mechanisms of skin wound healing. Nat. Rev. Mol. Cell Biol. 10.1038/s41580-024-00715-1 (2024).

3. M. T. Longaker et al., A Randomized Controlled Trial of the embrace Advanced Scar Therapy Device to Reduce Incisional Scar Formation. Plast. Reconstr. Surg. 134, 536–546 (2014).

4. H. I. Harn et al., The tension biology of wound healing. Exp. Dermatol. 28, 464–471 (2019).

5. C. K. Hsu et al., Mechanical forces in skin disorders. J. Dermatol. Sci. 90, 232–240 (2018).

6. G. C. Limandjaja, F. B. Niessen, R. J. Scheper, S. Gibbs, The Keloid Disorder: Heterogeneity, Histopathology, Mechanisms and Models. Front. Cell Dev. Biol. 8, 360 (2020).

7. R. Ogawa et al., The relationship between skin stretching/contraction and pathologic scarring: The important role of mechanical forces in keloid generation. Wound Repair Regen. 20, 149–157 (2012).

8. S. Akaishi, M. Akimoto, R. Ogawa, H. Hyakusoku, The Relationship Between Keloid Growth Pattern and Stretching Tension. Ann. Plast. Surg. 60, 445–451 (2008).

9. T. Nagasao et al., Transformation of keloids is determined by stress occurrence patterns on perikeloid regions in response to body movement. Med. Hypotheses 81, 136–141 (2013).

10. D. S. Foster et al., Integrated spatial multiomics reveals fibroblast fate during tissue repair. Proc. Natl. Acad. Sci. U.S.A. 118 (2021).

11. J. He et al., Mechanical stretch promotes hypertrophic scar formation through mechanically activated cation channel Piezo1. Cell Death Dis. 12, 226 (2021).

12. S. Mascharak et al., Preventing Engrailed-1 activation in fibroblasts yields wound regeneration without scarring. Science 372 (2021).

13. V. W. Wong et al., Focal adhesion kinase links mechanical force to skin fibrosis via inflammatory signaling. Nat. Med. 18, 148–152 (2011).

14. S. Zhou et al., A Novel Model for Cutaneous Wound Healing and Scarring in the Rat. Plast. Reconstr. Surg. 143, 468–477 (2019).

15. S. Aarabi et al., Mechanical load initiates hypertrophic scar formation through decreased cellular apoptosis. FASEB J. 21, 3250–3261 (2007).

16. N. Gao et al., Targeted inhibition of YAP/TAZ alters the biological behaviours of keloid fibroblasts. Exp. Dermatol. 10.1111/exd.14466 (2021).

17. Z. Jiang et al., Low-Frequency Ultrasound Sensitive Piezo1 Channels Regulate Keloid-Related Characteristics of Fibroblasts. Adv. Sci. 10.1002/advs.202305489 (2024).

18. C. C. Deng et al., Single-cell RNA-seq reveals fibroblast heterogeneity and increased mesenchymal fibroblasts in human fibrotic skin diseases. Nat. Commun. 12, 3709 (2021).

19. C. Feng et al., Single-cell RNA sequencing reveals distinct immunology profiles in human keloid. Front. Immunol. 13, 940645 (2022).

20. X. Liu et al., Single-Cell RNA-Sequencing Reveals Lineage-Specific Regulatory Changes of Fibroblasts and Vascular Endothelial Cells in Keloids. J. Invest. Dermatol. 142, 124–135 e111 (2022).

21. J. Shim et al., Integrated analysis of single-cell and spatial transcriptomics in keloids: Highlights on fibro-vascular interactions in keloid pathogenesis. J. Invest. Dermatol. 10.1016/j.jid.2022.01.017 (2022).

22. M. Direder et al., Schwann cells contribute to keloid formation. Matrix Biol. 108, 55–76 (2022).

23. Y. Xia, Y. Wang, M. Shan, Y. Hao, Z. Liang, Decoding the molecular landscape of keloids: new insights from single-cell transcriptomics. Burns & Trauma 11 (2023).

24. F. A. Wolf, P. Angerer, F. J. Theis, SCANPY: large-scale single-cell gene expression data analysis. Genome Biol. 19 (2018).

25. S. L. Wolock, R. Lopez, A. M. Klein, Scrublet: Computational Identification of Cell Doublets in Single-Cell Transcriptomic Data. Cell Syst. 8, 281–291.e289 (2019).

26. K. Polański et al., BBKNN: fast batch alignment of single cell transcriptomes. Bioinformatics 36, 964–965 (2020).

27. V. A. Traag, L. Waltman, N. J. van Eck, From Louvain to Leiden: guaranteeing well-connected communities. Sci. Rep. 9 (2019).

28. R. Satija, J. A. Farrell, D. Gennert, A. F. Schier, A. Regev, Spatial reconstruction of single-cell gene expression data. Nat. Biotechnol. 33, 495–502 (2015).

29. M. F. Griffin et al., JUN promotes hypertrophic skin scarring via CD36 in preclinical in vitro and in vivo models. Sci. Transl. Med. 13, 13 (2021).

30. M. L. Whitfield, L. K. George, G. D. Grant, C. M. Perou, Common markers of proliferation. Nat. Rev. Cancer 6, 99–106 (2006).

31. M. Bartoschek et al., Spatially and functionally distinct subclasses of breast cancer-associated fibroblasts revealed by single cell RNA sequencing. Nat. Commun. 9 (2018).

32. L. Cords et al., Cancer-associated fibroblast classification in single-cell and spatial proteomics data. Nat. Commun. 14 (2023).

33. G. C. Limandjaja et al., Reconstructed human keloid models show heterogeneity within keloid scars. Arch. Dermatol. Res. 310, 815–826 (2018).

34. B. Van de Sande et al., A scalable SCENIC workflow for single-cell gene regulatory network analysis. Nat. Protoc. 15, 2247–2276 (2020).

35. M. Downes, P. Koopman, SOX18 and the Transcriptional Regulation of Blood Vessel Development. Trends Cardiovasc. Med. 11, 318–324 (2001).

36. G. Zhang et al., Spi1 regulates the microglial/macrophage inflammatory response via the PI3K/AKT/mTOR signaling pathway after intracerebral hemorrhage. Neural Regen. Res. 19, 161–170 (2024).

37. X. L. Bai et al., Myocyte enhancer factor 2C regulation of hepatocellular carcinoma via vascular endothelial growth factor and Wnt/β-catenin signaling. Oncogene 34, 4089–4097 (2015).

38. G. Matrone et al., Fli1+ cells transcriptional analysis reveals an Lmo2–Prdm16 axis in angiogenesis. Proc. Natl. Acad. Sci. U.S.A. 118 (2021).

39. Graeme M. Birdsey et al., The Endothelial Transcription Factor ERG Promotes Vascular Stability and Growth through Wnt/β-Catenin Signaling. Dev. Cell 32, 82–96 (2015).

40. Z. Xu et al., Endothelial deletion of SHP2 suppresses tumor angiogenesis and promotes vascular normalization. Nat. Commun. 12 (2021).

41. S. Cioffi et al., Tbx1 regulates brain vascularization. Hum. Mol. Genet. 23, 78–89 (2014).

42. P. H. Reitsma et al., The Transcription Factor SOX18 Regulates the Expression of Matrix Metalloproteinase 7 and Guidance Molecules in Human Endothelial Cells. PLoS ONE 7 (2012).

43. J. Overman et al., Pharmacological targeting of the transcription factor SOX18 delays breast cancer in mice. eLife 6, e21221 (2017).

44. L. M. Treanor, S. Zhou, T. Lu, C. G. Mullighan, B. P. Sorrentino, Lmo2 Overexpression and Arf Loss Induce Myeloid Differentiation in Primitive Thymocytes. Blood 118, 47–47 (2011).

45. J.-N. Eckardt et al., Mutated IKZF1 is an independent marker of adverse risk in acute myeloid leukemia. Leukemia 37, 2395–2403 (2023).

46. S. J. Loughran et al., The transcription factor Erg is essential for definitive hematopoiesis and the function of adult hematopoietic stem cells. Nat. Immunol. 9, 810–819 (2008).

47. S. Stehling-Sun, J. Dade, S. L. Nutt, R. P. DeKoter, F. D. Camargo, Regulation of lymphoid versus myeloid fate ‘choice’ by the transcription factor Mef2c. Nat. Immunol. 10, 289–296 (2009).

48. A. Zakrzewska et al., Macrophage-specific gene functions in Spi1-directed innate immunity. Blood 116, e1-e11 (2010).

49. H. E. Talbott, S. Mascharak, M. Griffin, D. C. Wan, M. T. Longaker, Wound healing, fibroblast heterogeneity, and fibrosis. Cell Stem Cell 29, 1161–1180 (2022).

50. C. F. Guerrero-Juarez et al., Single-cell analysis reveals fibroblast heterogeneity and myeloid-derived adipocyte progenitors in murine skin wounds. Nat. Commun. 10, 650 (2019).

51. X.-M. Meng et al., Inflammatory macrophages can transdifferentiate into myofibroblasts during renal fibrosis. Cell Death Dis. 7, e2495–e2495 (2016).

52. S. Shen et al., Single-cell RNA sequencing reveals S100a9hi macrophages promote the transition from acute inflammation to fibrotic remodeling after myocardial ischemia=reperfusion. Theranostics 14, 1241–1259 (2024).

53. S. Eura et al., Hemodynamics and Vascular Histology of Keloid Tissues and Anatomy of Nearby Blood Vessels. Plast. Reconstr. Surg. Glob. Open 10, e4374 (2022).

54. S. SenGupta, C. A. Parent, J. E. Bear, The principles of directed cell migration. Nat. Rev. Mol. Cell Biol. 22, 529–547 (2021).

55. M. M. Nava et al., Heterochromatin-Driven Nuclear Softening Protects the Genome against Mechanical Stress-Induced Damage. Cell 181, 800–817 e822 (2020).

56. L. Kumar, M. E Futschik, Mfuzz: a software package for soft clustering of microarray data. Bioinformation 2, 5–7 (2007).

57. A. G. Solis et al., Mechanosensation of cyclical force by PIEZO1 is essential for innate immunity. Nature 573, 69–74 (2019).

58. Z. Wang et al., Endothelial upregulation of mechanosensitive channel Piezo1 in pulmonary hypertension. Am. J. Physiol. Cell Physiol. 321, C1010–C1027 (2021).

59. B. U, B. TW, Keloids: A Review of Etiology, Prevention, and Treatment. J. Clin. Aesthet. Dermatol. 13, 33–43 (2020).

60. N. C. Henderson, F. Rieder, T. A. Wynn, Fibrosis: from mechanisms to medicines. Nature 587, 555–566 (2020).

61. M. Vierhout et al., Monocyte and macrophage derived myofibroblasts: Is it fate? A review of the current evidence. Wound Repair Regen. 29, 548–562 (2021).

62. M. Aoki et al., The Immunosuppressant Fingolimod (FTY720) for the Treatment of Mechanical Force-Induced Abnormal Scars. J. Immunol. Res. 2020, 1–11 (2020).

63. R. Ogawa, The Most Current Algorithms for the Treatment and Prevention of Hypertrophic Scars and Keloids: A 2020 Update of the Algorithms Published 10 Years Ago. Plast. Reconstr. Surg. 149, 79e–94e (2021).

64. A. Leong, M. Kim, The Angiopoietin-2 and TIE Pathway as a Therapeutic Target for Enhancing Antiangiogenic Therapy and Immunotherapy in Patients with Advanced Cancer. Int. J. Mol. Sci. 21 (2020).

65. N. Rigamonti et al., Role of Angiopoietin-2 in Adaptive Tumor Resistance to VEGF Signaling Blockade. Cell Rep. 8, 696–706 (2014).

66. A. Scholz et al., Endothelial cell-derived angiopoietin-2 is a therapeutic target in treatment-naive and bevacizumab-resistant glioblastoma. EMBO Mol. Med. 8, 39–57 (2015).

67. M. Schmittnaegel et al., Dual angiopoietin-2 and VEGFA inhibition elicits antitumor immunity that is enhanced by PD-1 checkpoint blockade. Sci. Transl. Med. 9, eaak9670 (2017).

68. A. M. Khanani et al., The real-world efficacy and safety of faricimab in neovascular age-related macular degeneration: the TRUCKEE study – 6 month results. Eye 37, 3574–3581 (2023).

69. D. Lavie, A. Ben-Shmuel, N. Erez, R. Scherz-Shouval, Cancer-associated fibroblasts in the single-cell era. *Nat*. Cancer 3, 793–807 (2022).

70. L. T. S. Nguyen, M. A. C. Jacob, E. Parajón, D. N. Robinson, Cancer as a biophysical disease: Targeting the mechanical-adaptability program. Biophys. J. 121, 3573–3585 (2022).

