## Supplementary table 4 for "CD74^+^ fibroblasts proliferate upon mechanical stretching to promote angiogenesis in keloid"

***Corresponding author:**

**This PDF file includes:**

Supporting text

Tables S1-S3

**Materials and Methods**

**Analysis of single-cell sequencing data**

Analysis was performed on a rack-mount server with a Linux operating system (Red Hat 7.9). The following software and packages were used: anaconda (version 22.9.0), python (version 3.10.9), scanpy (version 1.9.3), anndata (version 0.9.1), bbknn(version 1.5.1), faiss (version 1.7.2), scvelo (version 0.2.5), pyscenic (version 0.12.0). The integration of datasets was done with the function annData.concatenate. Calculations of the mechanical score and the angiogenic score was done with the function scanpy.tl.score_genes.

**Cell culture**

Human foreskin fibroblasts (HFF, CTCC-001-0346) were isolated from human male foreskin biopsies. Keloid fibroblasts (KF, CTCC-185-HUM) were isolated from human keloid specimens. Both cell lines were cultured in Dulbecco’s MEM (C3113, Vivacell) supplemented with 10% fetal bovine serum (CTCC-002-001, Meisen) and 1% penicillin-streptomycin (Pen-Strep, C3420, Vivacell). Human umbilical vein endothelial cells (HUVEC, CTCC-003-0025) were cultured in endothelial cell complete medium (1001, ScienCell). All cells were purchased from Zhejiang Meisen Cell Technologies and were routinely PCR-tested for mycoplasma contamination (MP004, GeneCopoeia). All cells were cultured in a humidified incubator in 5% CO_2_ at 37 °C (Heracell 150i, Thermo Scientific).

**Manufacture of elastomer and cell stretching device**

The elastomer was made by casting polydimethylsiloxane (PDMS, Sylgard 184, Dow Corning) into polytetrafluoroethylene (PTFE) molds. Firstly, the base and curing agent (40:1 wt/wt) was weighted and vigorously mixed for 10 minutes in a paper cup. Then, the mixture was casted into the mold on an electronic scale to ensure consistent inter-elastomer thickness. Approximately 7 grams of the mixture is sufficient for the manufacture of one elastomer. Then, the mixture was degassed for 2 hours with a vacuum chamber, followed by curation at 65 °C for 18 hours. After carefully demolded, the elastomer was soaked in 75% ethanol for 15 minutes, air-dried and UV-sterilized on each side for 1 hour. The elastomer was coated with 30 µg/mL fibronectin solution for 1.5 hours at 37 °C and rinsed twice with DPBS. The cell attachment area of the elastomer is 20*20 mm^2^. The cell stretching device was assembled by mounting a DC motor (TS-DIY-37GB330, Tsiny Motor), a pair of linear guiding rails (SBR12, Shunhe Shengda Automation Technologies), and a pair of 3D-printed elastomer holders to a PTFE baseplate with dimensions of 300*210*4 mm^3^. The motor was actuated by a 15-volt DC power supply. The model file of the elastomer mold and the 3D-printed elastomer holder is available with the link <https://www.tinkercad.com/things/eFT8PcZPDnA-stretching-device-for-incubator?sharecode=vdXOSe14MuX6g8tGPGuVTep4Ie_GPVvI2cAiykPY-1M>.

**Mechanical stretching of cultured cells**

Before stretching experiments, 8*10^4^ cells were seeded into the elastomer and were allowed to adhere for 24 hours. Then, the elastomer was installed on the cell stretching device and cells were stretched for the indicated time at a frequency of 0.1 Hz in an incubator. The maximum elongation of the elastomer was 50% of its resting length.

**BrdU labelling and immunofluorescence of cells**

For BrdU labelling of cells, cells were incubated in 10 µM BrdU (B5002, Sigma) for 12 hours before fixation. The elastomer was cut with an 8-mm circular cutting die into four round slices. Then, cells on the slice were fixed in 4% paraformaldehyde (PFA, BL539A, Biosharp) for 15 minutes, permeabilized with 0.25% Triton X-100 in DPBS for 10 minutes, hydrolyzed with 1.5 M HCl for 20 minutes, and blocked in 5% bovine serum albumin (BSA, 4240GR005, Biofroxx) at room temperature for 1 hour. Cells were subsequently incubated overnight in primary antibody diluted in 5% BSA/0.25% TritonX-100/DPBS at a 4 °C cold room, followed by washing of three times in DPBS and incubation in secondary antibody at room temperature for 1 hour. Finally, samples were stained with DAPI for 10 minutes, washed three times and mounted on a rectangular coverslip (CLS2975224, Corning) with Diamond antifade mountant (S36963, Invitrogen). The following antibodies and reagents were used: Alexa Fluor 647 Phalloidin (1:300, A22287, Invitrogen), BrdU (1:300, 555627, BD), and DAPI (1 µg/mL, D9542, Sigma).

**Cryosection and immunofluorescence of keloid specimen**

The fresh keloid specimen was fixed in 4% PFA for 24 hours, placed in 30% sucrose at 4 °C until sink, embedded in Optimal Cutting Temperature (OCT, 4583, Sakura Finetek), and cryosectioned into 15-µm thickness with a cryotome (CM1950, Leica). Then, slides were subject to antigen retrieval in pH 9 antigen retrieval buffer at 98 °C for 20 minutes, blocked in 5% BSA for 1 hour, and stained overnight in a cold room with anti-PDGFRA antibody. Slides were then rinsed three times, stained with Alexa Fluor 647-secondary antibody for 1 hour, rinsed three times, stained overnight in a cold room with fluorophore-conjugated CD31 and CD74 antibodies, rinsed three times, stained with DAPI for 10 minutes, rinsed, and mounted. The following antibodies and reagents were used: PDGFRA (1:200, 3174, CST), Alexa Fluor 647-Goat anti rabbit secondary antibody (1:300, A21244, Invitrogen), CoraLite Plus 488-CD31 (1:100, CL488-11265, Proteintech), PE-CD74 (1:20, E-AB-F1072D, Elabscience), and DAPI (1 µg/mL, D9542, Sigma).

**Confocal microscopy and image analysis**

All fluorescence images were collected on a spinning-disk confocal microscope (CSU-W1, Nikon). Images were acquired as confocal stacks and projected along Z axis to obtain maximum intensity projection images. Same image acquisition settings were used for all samples within the same set of experiment. Images were analyzed using NIS-Elements AR software (version 5.42.02, Nikon). The function ‘define threshold’ was used for cell segmentation, and the function ‘binary layers’ was used to select cells positive for more than one marker.

**RNA sequencing and data analysis**

Total RNA was isolated using RNAiso plus (9109, Takara) and quantified using Nanodrop One. The quality of RNA was determined by Bioanalyzer 2100 (Agilent Technologies). After the RNA was purified by poly-T magnetic beads, sequencing libraries was generated, quantified, and sequenced on a NovaSeq 6000 sequencer (Illumina) with PE150 strategy. For RNA-seq data analysis, raw fastq files were processed with trim_galore (version 0.6.7), mapped with STAR (version 2.7.10b) and quantified with featurecounts (version 2.0.5) in Linux (Red Hat 7.9). The GRCh38 reference genome (version p13) was used for mapping and GENCODE (version 32) annotation file was used for quantification. Differential expression analysis was performed in R (version 4.2.1) using DESeq2 (version 1.38.3). Functional enrichment analysis was performed using clusterprofiler (version 4.6.2). Heatmap was generated with pheatmap (version 1.0.12). GSVA analysis was performed with GSVA (version 1.46.0).

**HUVEC tube formation assay**

Matrigel (356231, Corning) with a volume of 40 µL was added to the well of a 96-well cell culture plate, and the plate was centrifuged at 3000g for 30 minutes to even the surface of the gel. Then, 7*10^3^ HUVEC cells were suspended in the culture media indicated and plated into the well. After 4 hours, cells were stained with calcein-AM (C2012, Beyotime) and imaged under a microscope. The image quantification was done by AngioTool (version 0.6a, NIH).

**Transwell migration assay**

Before the assay, cells were serum-starved in 0.5% FBS/DMEM for 24 hours. Then, 2.5*10^5^ cells were suspended in 200 µL 0.5% FBS/DMEM and seeded into the transwell chamber with 8-μm pores (3422, Corning). 10% FBS/DMEM were added into the lower chamber as chemoattractant. After incubation of 24 hours, cells were fixed with 4% PFA for 10 minutes, and cells remaining in the upper chamber were removed with a wetted cotton swab. Then, the chamber was air-dried, stained with crystal violet for 10 minutes, and washed with deionized water. After the chamber was air-dried, the membrane was removed from the chamber with a 11# scalpel and mounted on a glass slide with neutral balsam. Then, bright-field images were acquired with a microscope.

**Quantitative RT-PCR**

Total RNA with a quantity of 1 µg was reverse-transcribed into cDNA (K16225, Thermo) and aliquoted. Then, 20 ng of cDNA was quantified with quantitative PCR using SYBR green (QST-100, Toroivd) on a QPCR instrument (Quantstudio 3, Applied Biosystems). Sequences of primers used in this study were: PIEZO1-F: ATGTTGCTCTACACCCTGACC; PIEZO1-R: CCAGCACACACATAGATCCAGT; GAPDH-F: GTCTCCTCTGACTTCAACAGCG, GAPDH-R: ACCACCCTGTTGCTGTAGCCAA.

**Fluorescence-activated cell sorting (FACS)**

HFF and KF cells were trypsinized, filtered through a 70 µm nylon cell strainer (15-1070, Biologix), labelled with CD74-PE (1:20, E-AB-F1072D, Elabscience) for 30 minutes at 4 °C, and sorted by a FACSAria IIu (BD) cell sorter. Gates were initially established on forward scatter and side scatter to select all cells. Then, staining of PE-Mouse IgG1 Isotype control (1:20, E-AB-F09792D, Elabscience) was used to determine appropriate gating for CD74+ fibroblasts. Sorted cells were collected in 15-mL falcon tubes containing 5% Pen-Strep/20% FBS/DMEM.

**Transfection and RNAi**

siRNA sequences targeting hPiezo1 were: siPIEZO1-1: CCAAGUACUGGAUCUAUGU(dT)(dT), siPIEZO1-2: CCAAGAAGUACAAUCAUCU(dT)(dT), siNC: UUCUCCGAACGUGUCACGU (dT)(dT). Transfections were performed using Lipofectamine 3000 (Invitrogen) and opti-MEM (Gibco) according to manufacturer’s instructions. Cells were subjected to the experimental analysis indicated after 48 hours of transfection.

**Western blotting**

Cells were rinsed in DPBS and lysed in 2% SDS lysis buffer supplemented with protease (P8340, Sigma) and phosphatase inhibitors (CW2383S, CWBio). After sonication, samples were quantified using BCA assay (P0009, Beyotime). The cell lysate was then reduced in Laemmli sample buffer at 95 °C for 5 minutes and resolved by SDS-PAGE gels and transferred onto nitrocellulose membranes (66485, Pall). Membranes were blocked with 5 % non-fat milk in tris-buffered saline containing 0.1% tween-20 (TBST) for 1 hour at room temperature, after which diluted primary antibodies were added and incubated over night at 4 °C on a shaker. The primary antibody was diluted with 5% BSA/TBST. The membranes were subsequently washed in TBST for three times, and incubated in secondary HRP-conjugated antibodies (31430 and 31460, Invitrogen) diluted with 5% milk/TBST for 1 hour at room temperature. After extensive washing in TBST, antibody binding was detected by immobilon western HRP substrate (WBKLS0500, Millipore) using a chemiluminescence imaging system and the images were quantified using FIJI (version 2.9.0, NIH). The following antibodies were used: Phospho-AKT (Ser473) (1:1000, 66444-1-lg, Proteintech), AKT (1:1000, 10176-2-AP, Proteintech), Phospho- ERK1/2 (Thr202/Tyr204) (1:1000, 28733-1-AP, Proteintech), ERK1/2(1:1000, 11257-1-AP, Proteintech), GAPDH (1:3000, GB11002, Servicebio).

**Calcium imaging of mechanically stretched cells**

For the manufacture of the stage-top stretching device, two linear actuators (MS2006-48, Ying Pengfei) and two microswitches were mounted to a 3D-printed enclosure. The actuator was connected to and powered by a stepping motor drive (MD-2422, Samsr motor) under the control of a single-axis motion controller (HJ-03A, Haijie Jiachuang Technologies). The model file of the 3D-printed enclosure is available with the link <https://www.tinkercad.com/things/jE9T7bTF43e-stage-top-stretcher?sharecode=A7tfdiNYUdckl4hBCTWQIxxSwUBpOqs4cQVrHTcSnj0>. For imaging of stretched cells, 8*10^4^ cells were seeded to the PDMS elastomer and allowed to adhere for 24 hours. Then, cells were stained with Fluo-4 AM (S1061S, Beyotime) and Hoechst 33342 (C1022, Beyotime) for 30 minutes, followed by washing with fresh media. Then, the elastomer was mounted to the stage-top stretching device, and the device was placed on the microscope stage equipped with incubation chamber (Tokai Hit). Then, images were taken to establish the baseline fluorescence signal intensity, after which cells were stretched for 20 cycles, and imaged at indicated timepoints.

**Supplementary Table 1. Patient demographics of all samples across 5 datasets**

| Accession | Sample | Sex | Age | Location |
| --- | --- | --- | --- | --- |
| GEO-  GSE163973 | Dataset 1 keloid 1 | M | 20 | Back |
|  | Dataset 1 keloid 2 | M | 23 | Chest |
|  | Dataset 1 keloid 3 | F | 34 | Chest |
|  | Dataset 1 normal scar 1 | M | 39 | Back |
|  | Dataset 1 normal scar 2 | M | 28 | Chest |
|  | Dataset 1 normal scar 3 | F | 26 | Chest |
| GSA –  HRA000425 | Dataset 2 keloid 1 | F | 28 | Chest |
|  | Dataset 2 keloid 2 | F | 32 | Chest |
|  | Dataset 2 keloid 3 | F | 26 | Chest |
|  | Dataset 2 keloid 4 | F | 26 | Chest |
|  | Dataset 2 skin 1 | F | 28 | Chest |
|  | Dataset 2 skin 2 | F | 32 | Chest |
|  | Dataset 2 skin 3 | F | 26 | Chest |
|  | Dataset 2 skin 4 | F | 26 | Chest |
| GEO –  GSE181297 | Dataset 3 keloid 1 | M | 23 | Ear |
|  | Dataset 3 keloid 2 | M | 25 | Back |
|  | Dataset 3 normal scar 1 | M | 25 | Back |
| GEO –  GSE181316 | Dataset 4 keloid 1 | F | 36 | Sternum |
|  | Dataset 4 keloid 2 | F | 60 | Earlobe |
|  | Dataset 4 keloid 3L | M | 34 | Earlobe (Left) |
|  | Dataset 4 keloid 3R | M | 34 | Earlobe (Right) |
|  | Dataset 4 normal scar 1 | F | 26 | Abdomen |
|  | Dataset 4 normal scar 2 | F | 43 | Abdomen |
|  | Dataset 4 normal scar 3 | M | 60 | Abdomen |
|  | Dataset 4 skin 1 | M | 33 | Abdomen |
| GEO-  GSE130973 | Dataset 5 Skin 1 | M | 25 | Inguinal region |
|  | Dataset 5 Skin 2 | M | 27 | inguinal region |
|  | Dataset 5 Skin 3 | M | 53 | inguinal region |
|  | Dataset 5 Skin 4 | M | 70 | inguinal region |
|  | Dataset 5 Skin 5 | M | 69 | inguinal region |

**Supplementary Table 2. Quantity and relative abundance of each cell type among all samples**

| Cell type | Groups | | | |
| --- | --- | --- | --- | --- |
|  | Keloid | Scar | Skin | All samples |
| Endothelial cell | 29379 (26.3%) | 7880 (17.4%) | 6229 (11.5%) | 43488 (20.6%) |
| Erythrocyte | 111 (0.1%) | 189 (0.4%) | 53 (0.1%) | 353 (0.2%) |
| Fibroblast | 42476 (38.1%) | 16868 (37.3%) | 13423 (24.9%) | 72767 (34.5%) |
| Keratinocyte | 13836 (12.4%) | 10180 (22.5%) | 17829 (33.0%) | 41845 (19.9%) |
| Lymphatic cell | 3089 (2.8%) | 1474 (3.3%) | 5101 (9.4%) | 9664 (4.6%) |
| Lymphatic EC | 2680 (2.4%) | 1049 (2.3%) | 832 (1.5%) | 4561 (2.2%) |
| Macrophage | 3810 (3.4%) | 2600 (5.7%) | 4433 (8.2%) | 10843 (5.1%) |
| Mast cell | 1779 (1.6%) | 277 (0.6%) | 394 (0.7%) | 2450 (1.2%) |
| Melanocyte | 127 (0.1%) | 466 (1.0%) | 624 (1.2%) | 1217 (0.6%) |
| Neural cell | 1674 (1.5%) | 107 (0.2%) | 399 (0.7%) | 2180 (1.0%) |
| Mural cell | 11848 (10.6%) | 4076 (9.0%) | 4283 (7.9%) | 20207 (9.6%) |
| Sweat gland cell | 725 (0.7%) | 60 (0.1%) | 407 (0.8%) | 1192 (0.6%) |
| Total | 111534 | 45226 | 54007 | 210767 |

**Supplementary Table 3.** **Quantity and relative abundance of each fibroblast subtype among all samples**

| Cell type | Groups | | | |
| --- | --- | --- | --- | --- |
|  | Keloid | Scar | Skin | All samples |
| Cluster 0 | 3787 (8.9%) | 651 (3.9%) | 304 (2.3%) | 4742 (6.5%) |
| Cluster 1 | 2492 (5.9%) | 177 (1.0%) | 68 (0.5%) | 2737 (3.8%) |
| Cluster 2 | 15525 (36.6%) | 482 (2.9%) | 170 (1.3%) | 16177 (22.2%) |
| Cluster 3 | 6737 (15.9%) | 254 (1.5%) | 106 (0.8%) | 7097 (9.8%) |
| Cluster 4 | 3942 (9.3%) | 5431 (32.2%) | 3177 (23.7%) | 12550 (17.2%) |
| Cluster 5 | 314 (0.7%) | 773 (4.6%) | 1073 (8.0%) | 2160 (3.0%) |
| Cluster 6 | 790 (1.9%) | 374 (2.2%) | 180 (1.3%) | 1344 (1.8%) |
| Cluster 7 | 244 (0.6%) | 219 (1.3%) | 157 (1.2%) | 620 (0.9%) |
| Cluster 8 | 1312 (3.1%) | 1451 (8.6%) | 2885 (21.5%) | 5648 (7.8%) |
| Cluster 9 | 5441 (12.8%) | 3852 (22.8%) | 2295 (17.1%) | 11588 (15.9%) |
| Cluster 10 | 1892 (4.5%) | 3204 (19.0%) | 3008 (22.4%) | 8104 (11.1%) |
| Total | 42476 | 16868 | 13423 | 72767 |

**Supplementary Table 4. Result of differential gene expression analysis of fibroblast subtype 0 versus fibroblast subtypes 1, 2, and 3 from all samples.**
